# *Phaeocystis* blooms caused carbon drawdown during the Antarctic Cold Reversal from sedimentary ancient DNA

**DOI:** 10.1101/2024.04.11.589015

**Authors:** Josefine Friederike Weiß, Ulrike Herzschuh, Juliane Müller, Jie Liang, Maria-Elena Vorrath, Amedea Perfumo, Kathleen R. Stoof-Leichsenring

**Author notes:** Correspondence author: Kathleen R. Stoof-Leichsenring. Shared senior authorship (Ulrike Herzschuh, Kathleen R. Stoof-Leichsenring).

## Abstract

The Southern Ocean plays a crucial role in the global carbon budget. Modeling studies propose that the atmospheric CO_2_ plateau during the Antarctic Cold Reversal (ACR; 14,700 to 12,700 calibrated years before present (cal yr BP)) is related to increased marine productivity. However, proxy evidence relating environmental conditions as well as primary community composition and productivity to carbon drawdown is missing. Our ancient DNA shotgun metagenomic analysis of marine sediments revealed *Phaeocystis antarctica* (haptophyte) as a key element of the primary producer community. Independent proxy evidence (blooming-related bacteria, Ba/Fe ratio) from the same sediment record point to high productivity in response to enhanced sea-ice seasonality caused by ACR cooling. Post ACR, abrupt *Phaeocystis* community loss shows how sensitive this ecosystem is to warming, potentially representing a key tipping element that may be further enhanced by the *Phaeocystis*-related sulfur cycle–climate feedback. As an analogy for present warming, it highlights the importance of regions with high seasonal sea-ice variability and *Phaeocystis*-dominance, such as the Ross Sea, for stabilizing atmospheric CO_2_ content. Additionally, our shotgun metagenomic data portray complex Holocene ecosystem establishment including key Antarctic taxa such as penguins, whales, and Antarctic fishes with implications for ongoing conservation efforts.

## Main

The Southern Ocean, surrounding the Antarctic continent, plays a key role in the global carbon budget (1). It has been estimated to absorb more than 40% of the total CO_2_ emitted due to human activity since the start of industrialization (2). Marine phytoplankton are the most important primary producers in the ocean and account for a large portion of CO_2_ drawdown (3). Carbon dioxide assimilated by phytoplankton is exported in the form of organic carbon to deeper water layers (drawdown) and, at least partly, sequestered to the sediments (export). This so-called biological carbon pump slows down carbon remineralization and re-emission and operates as an important long-term carbon sink (4). It is not yet understood how strongly global warming will impact the effectiveness of the biological carbon pump in Antarctica (5) especially with the loss of sea ice (6).

The global warming at the end of the last glacial period (21,000–11,700 cal yr BP) was interrupted by a short cold period in the Southern Hemisphere, termed the Antarctic Cold Reversal (ACR; 14,700–12,700 cal yr BP) (7). A previous study has shown that the sea-ice extent varied strongly between the seasons during the ACR, meaning that there was a large difference between summer and winter sea-ice areas (8). It has been speculated, based on modeling, that the resulting high productivity and reduced outgassing facilitated a global CO_2_ plateau and thus initiated a dampening feedback loop on global temperature during the ACR (8). However, the paleo evidence is uncertain, mainly because we lack detailed proxy knowledge about past carbon drawdown and export during the ACR that may have led to a plateau in atmospheric CO_2_ concentrations (9). Biogenic opal concentration in sediments is the most commonly used paleoproductivity and carbon export proxy (10). Studies have found a decline or stable low concentration of biogenic opal during the ACR (11), which is inconsistent with the assumption that diatom blooms in the Southern Ocean were of critical importance in the ACR CO_2_ plateau (8). High Ba/Fe ratios may indicate organic carbon drawdown in the Southern Ocean (12) as barite (BaSO_4_) can precipitate during phytoplankton blooms, which is linked to carbon precipitation (13), but records covering the ACR are lacking.

The composition of the primary producer community plays a major role in the speed and effectiveness of carbon drawdown and export (14) because the productivity rates of the taxa differ (15). Sea-ice conditions, in turn, determine the composition of the phytoplankton community (16) in the Southern Ocean (17). Therefore, time series which place phytoplankton compositional turnover in the context of changing carbon drawdown and export, sea-ice conditions, and atmospheric CO_2_ are urgently needed.

The Antarctic Peninsula is known to be the fastest warming region of the Antarctic continent (18). Nowadays, the marine ecosystem in this region is mainly influenced by open ocean conditions (3). The primary producer community consists mainly of taxa associated with the open ocean (19), such as centric diatoms (e.g., *Chaetoceros*) (20) and Antarctic krill (21), which is the main food source for top predators like whales and penguins (22). Due to gaps in the fossil record for many taxa, it remains difficult to state since when they occurred in the ecosystem.

In the last two decades, sedimentary ancient DNA, a promising new paleoecological proxy, has been established. This cutting-edge method, which analyzes fragmented DNA strands bound to mineral particles in terrestrial and marine sediments, enables the characterization of biological community changes through time independently of fossil remains (23). The sedimentary ancient DNA (sedaDNA) metagenomic method is a non-target approach that bypasses the biases introduced by PCR amplification (24), known from metabarcoding (25). It allows the study of past communities at the ecosystem-level because it facilitates the detection of flora, fauna, and microorganisms concurrently (26). There has been one recent study that used an ecosystem-level metagenomic approach on marine sediments in the Bering Sea region (27).

For this study, we examined a sediment core from the northern Antarctic Peninsula region, specifically the Bransfield Strait (PS97/072-1; Fig. 1), spanning the last 14,000 years. Using sedaDNA shotgun metagenomics, we identified *Phaeocystis antarctica* as the dominant primary producer during the ACR in the high latitude Southern Ocean. By employing conventional proxies, such as the XRF derived Ba/Fe ratio, we were able to detect the increase in carbon drawdown linked to elevated *Phaeocystis antarctica* abundance during the ACR, which is additionally supported by a strong bacterioplankton blooming. Our findings reveal that as warming and a decrease in sea ice occurred, the modern ecosystem gradually established with the dominance of *Chaetoceros* at the end of the ACR.

**Figure 1:**
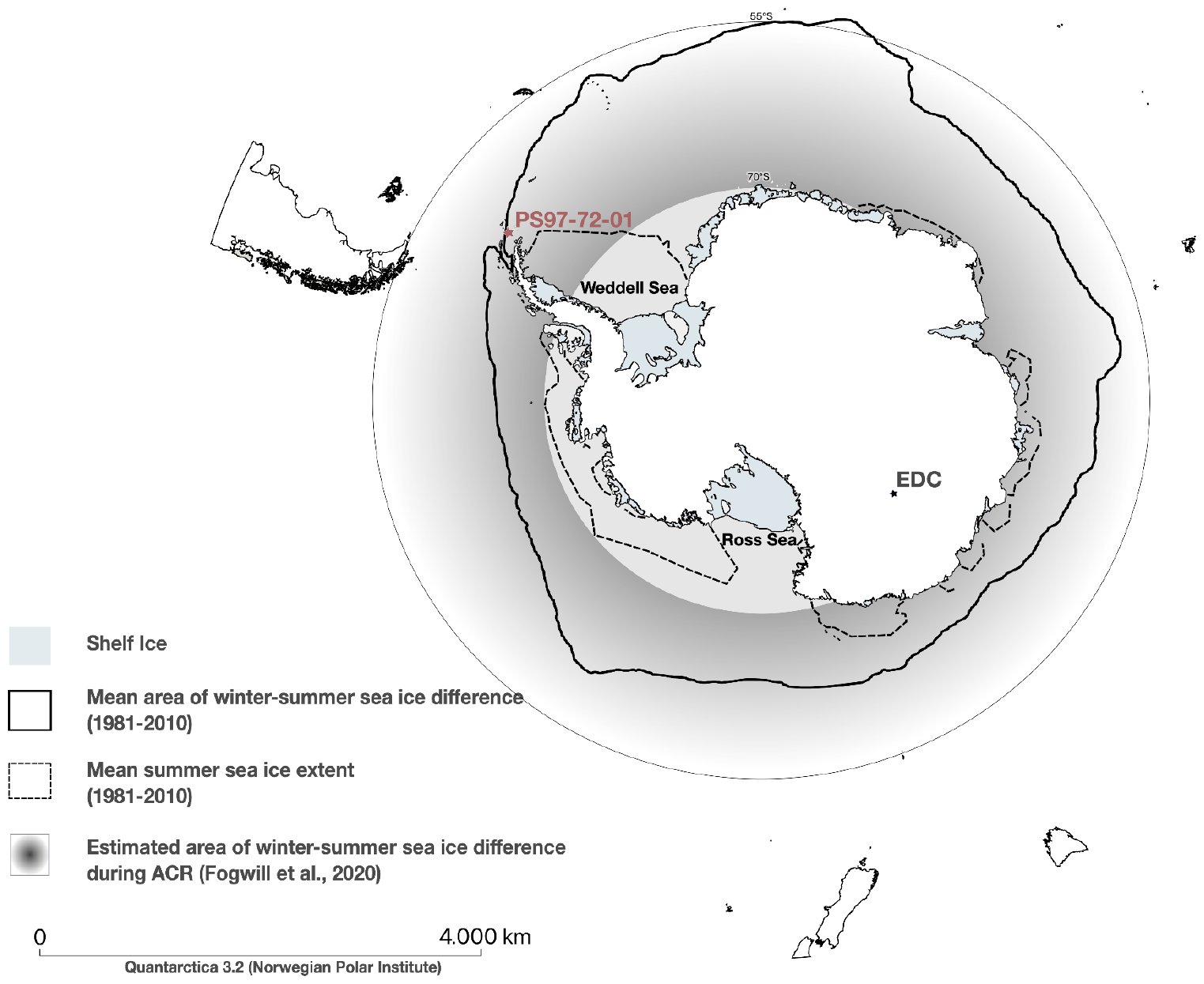
Seasonal difference of sea-ice area over the time periods of the Antarctic Cold Reversal (ACR: 14.7–12.7 k yrs BP) and the mean of the last 20 years around Antarctica with the location of the studied marine sediment core PS97/072-1 and the ice core EPICA Dome C (EDC). The mean area (approximately 15 million km^2^) between 1981-2010 of winter-summer sea-ice difference is shown in light gray (49). The modeled difference between winter sea-ice area (sea-ice extent up to 55°S in winter) and summer sea-ice area (sea-ice extent up to 70°S) during ACR is shown as a gradient gray area and reaches up to 20 million km^2^ (8). The Antarctic Polar Stereographic basemap was created using the SCAR Antarctic Digital Database version 7.0 in the QGis project Quantarctica 3.2 (Norwegian Polar Institute).

## Results & Discussion

### Phaeocystis antarctica dominates the phytoplankton community during the ACR

A total of 2,242,941 reads, retrieved from shotgun sequencing, were assigned to eukaryotic organisms including taxa from most of the major marine phyla (Bacillariophyta, Haptophyta, Chlorophyta, Chordata, Arthropoda) (Supplementary data 1,2). Our dataset is among the first marine datasets to allow an ecosystem-level reconstruction, and is unique for the Southern Ocean in having high resolution covering the ACR and the transition to the Holocene.

Phytoplankton reads dominate the dataset (1,952,627; 87%). The haptophyte *Phaeocystis antarctica* dominates the photoautotrophic community during the ACR comprising over 60%. It is accompanied by pennate diatoms of the genus *Fragilariopsis* (e.g., *Fragilariopsis cylindrus*), accounting for more than ∼15% of the total community (Fig. 2). Additionally, we identify Chlorophyta such as *Micromonas* as primary producers at the end of the ACR with a relative abundance of approximately 10%.

**Figure 2:**
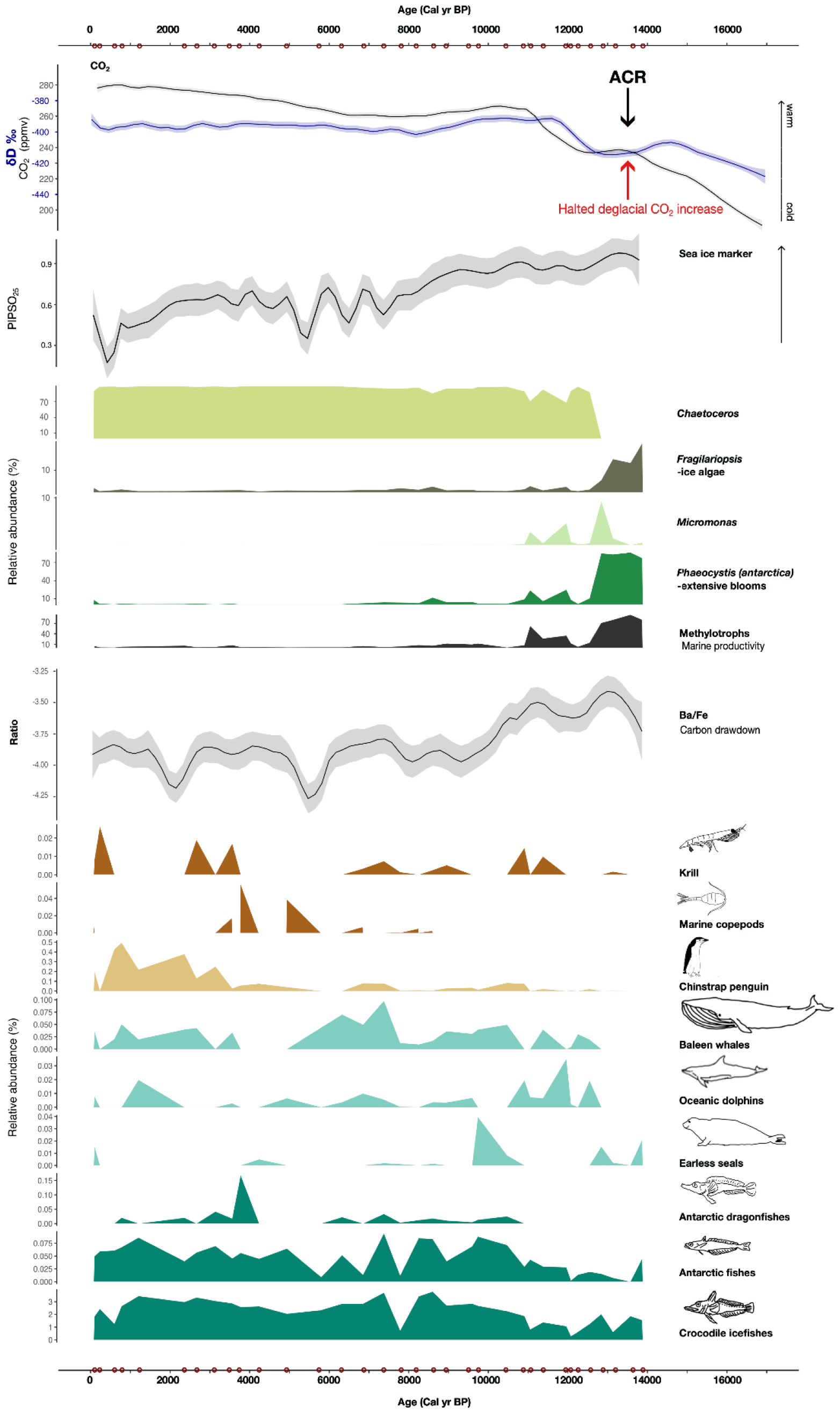
Primary producers and higher trophic level community composition over time. The upper most graph shows the δD ‰ (29) values for temperature reconstructions and the reconstructed atmospheric CO_2_ concentrations of the EPICA Dome C ice core for the last 17,000 years (50). The black arrow indicates the Antarctic Cold Reversal and the red arrow marks the halted deglacial CO_2_ rise. Following is the P_B_IPSO_25_ sea-ice biomarker graph over the last 14,000 years derived from the PS97/72-01 core (11). Next are the dominant primary producer taxa derived from shotgun metagenomics of sedaDNA over the last 14,000 years. In addition, the relative abundance of methylotrophic bacteria is shown, as well as the Ba/Fe ratio derived from elemental scanning. Finally, reconstructed ecosystem community composition of higher trophic levels including only Antarctic taxa over the last 14,000 years. The red dots on the x-axis show the discrete sampling points for sedaDNA shotgun metagenomics.

By analogy with present-day conditions this primary producer community is associated with ecosystems with higher sea-ice extent (16), such as those characterizing the present Ross Sea (28). The ACR was a period of strong cooling in the Southern Hemisphere during deglacial warming, which is reflected in the EPICA Dome C ice core (29). Biomarker-based sea-ice reconstruction, the higher occurrence of sea-ice diatoms in the fossil record of core PS97/072-1 (11), and the estimated winter-summer sea-ice difference based on model results (Fig. 1) confirm a spatial and temporal increase in sea-ice cover during the ACR (Fig. 2).

The main pattern in the photoautotrophic community in this study is a shift in dominance from a haptophyte-to a diatom-dominated community (Fig. 2) at approximately 12,700 cal yr BP at the end of the ACR. This shift between communities and the time point of change is supported by a Latent Dirichlet allocation (LDA) and a Bayesian change-point model (Supplementary Figure 1). The increase in temperature from ice-core data (29), the decline in sea-ice biomarkers (11), and the decrease in abundance of sea ice-associated taxa are congruent with the onset of a warm period beginning with the termination of the ACR (Fig. 2). Throughout the Holocene, the community is dominated by *Chaetoceros* species. The shift from *Phaeocystis* (haptophyte) to a diatom-dominated community at the end of the ACR is consistent with independent quantitative data on diatom concentration and biogenic opal (Supplementary Figure 2, Supplementary Table 1), which show low productivity and concentration of diatoms during the ACR and an increase in these values during the Holocene. Furthermore, qualitative changes in the composition can be seen in the independent morphological count data, which correlate significantly with the diatom data of the paleogenetic record (Supplementary Table 1). Additionally, this shift is supported by a study from the Scotia Sea showing an increased abundance of Chaetocerotaceae since 12,700 cal yr BP (26).

### Association of carbon drawdown with *Phaeocystis antarctica* during the Antarctic Cold Reversal

We have substantial paleo evidence that the reconstructed *Phaeocystis*-dominated sea-ice community in the Southern Ocean during the ACR was accompanied by high carbon drawdown. Recent *Phaeocystis*-dominated communities are characterized by biomass-rich blooming owing to their strategy to form colonies to reduce zooplankton grazing pressure (30). Moreover, *Phaeocystis antarctica* has significantly higher carbon dioxide absorption compared to diatoms and can achieve a near-maximum photosynthetic rate at low irradiances such as under sea ice (17).

Strong phytoplankton blooming during the ACR is accompanied by the bacterioplankton signal in our core. We observe that the period of high *Phaeocystis* abundance has a high co-occurrence of methylotrophic bacteria specializing in single carbon compound metabolism (Methylophilaceae, Methylobacteriaceae, Hyphomicrobiaceae, Beijerinckiaceae, Methylococcaceae, and *Methylophaga* (or their relatives)) in our prokaryote record. These known associations of algal blooms in modern oceans use methanol produced during the blooms as an energy source (31). From this we suggest that indications of methylotrophic bacteria in the shotgun metagenomics record may be used in the future, after further testing, as a new paleo proxy for past algal blooms.

The Ba/Fe ratio is a known proxy for carbon drawdown and export productivity, because barite (BaSO_4_) precipitates during phytoplankton blooms (32) (Supplementary data 3). From element scanning of our sediment core we observe a high Ba/Fe ratio during the *Phaeocystis*-rich ACR (Fig. 2). Against our expectations, recorded total organic carbon (TOC) values of the same core do not show increased values during the ACR (11). We speculate that captured carbon circulated in the deep sea and was not stored on longer time scales in the sediments. Because of the shortness of the ACR and subsequent warming, increased upwelling occurred, presumably leading to a strong outgassing of CO_2_ from the deeper layers (33) which, at least partly, may have caused the early Holocene CO_2_ rebound effect (34).

To test our hypothesis, we conducted a statistically temporal correlation analysis using a piecewise structural equation model (SEM), as shown in Figure 3A. The SEM method can effectively elucidate the complex relationships between paleogenetic data and various environmental parameters (35). We employed the SEM method to assess the direct and indirect influences of environmental factors and phytoplankton on the carbon cycle. We evaluated the model fit using Fisher’s C value (6.80) and the Chi-squared statistic (6.896), which assesses all directed separation tests. Model selection based on the Akaike information criterion (AIC) confirmed that our model, considering specific environmental factors and phytoplankton abundance, provided the best prediction of the carbon-cycle dynamics (Supplementary data 4). In particular, our results show a highly significant negative correlation of the temperature proxy (EDC δD ‰) and a highly significant positive correlation of the sea-ice proxy (PIPSO_25_) with the dominant primary producers during the ACR, *Phaeocystis*, and *Fragilariopsis*. In contrast, the opposite pattern emerges for the diatom *Chaetoceros*. Among the proxies reconstructing the carbon cycle, *Phaeocystis* shows a significant negative correlation with the reconstructed CO_2_ concentrations and conversely a significant positive correlation with the carbon drawdown proxy Ba/Fe ratio, which in turn, shows a significant inverse relationship with CO_2_ concentrations. In addition, *Phaeocystis* exhibits a significant correlation with methylotrophic bacteria as they increase in abundance due to a high biomass of primary producers. The methylotrophic bacterioplankton in turn exhibit the same relationship with the Ba/Fe ratio. In contrast, *Chaetoceros* exclusively exhibits a significant negative correlation with methylotrophic bacterioplankton. *Fragilariopsis* shows no significant correlation with the environmental parameters used to reconstruct the carbon cycle.

**Figure 3:**
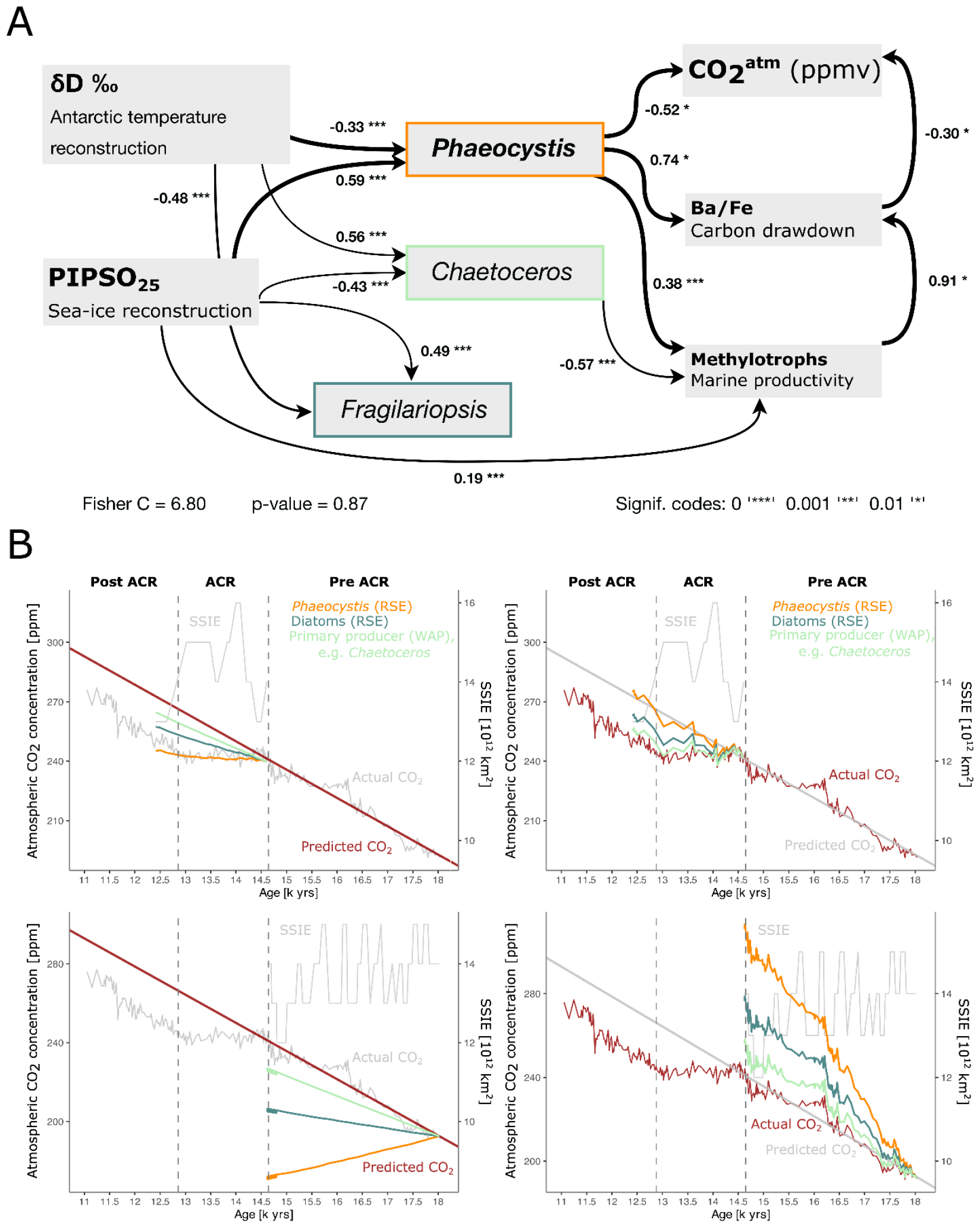
A Piecewise Structural Equation Model (SEM) of environmental parameters and paleogenetic data of the main primary producers. The goodness-of-fit of the piecewise SEM is presented as Fisher C = 6.80 and the overall model p-value = 0.87. **B Seasonal sea ice extent (SSIE) [10**^**12**^ **km**^**2**^**] (8) and calculation of the impact of net primary productivity (NPP) of different primary producers in the Southern Ocean in diverse environments (RSE = Ross Sea environment; WAP = Western Antarctic Peninsula) on global atmospheric carbon dioxide concentration [ppm] before, after and during the Antarctic cold reversal**. The left-hand fields show in red the extrapolation of the actual CO_2_ concentration from this period from which the NPP was subtracted. On the right-hand fields the real CO_2_ curve is shown in red and the NPP was added.

Once the temporal correlation was found to be significant we want to quantitatively assess our hypothesis, that a shift of diatom-dominance to *Phaeocystis antarctica* dominance in the increased seasonal sea-ice area resulted in the reconstructed global atmospheric carbon dioxide plateau and related ACR (Fig. 3B). For this purpose, we collected information about the influence of different major primary producers (*Phaeocystis antarctica* and diatoms) under different ecosystem conditions (modern Ross Sea environment and modern West Antarctic Peninsula environment) on the global atmospheric carbon dioxide concentration from the literature. In the Ross Sea, *Phaeocystis antarctica* shows an average net primary production (NPP) of 77.5 g C m^-2^ yr^-1^, while sea-ice diatoms like *Fragilariopsis* exhibit 38.9 g C m^-2^ yr^-1^ (36); in contrast, the Western Antarctic Peninsula region, dominated by diatoms like *Chaetoceros*, has a lower primary productivity of 15.83 g C m^-2^ yr^-1^ (37).

Using these values and the information about the dynamic sea-ice extent the drawdown of CO_2_ due to community-specific primary productions would accumulate up to 20.64 ppm (*Phaeocystis*), 10.36 ppm (sea ice diatoms, e.g. *Fragilariopsis*) or respectively 4.23 ppm (modern WAP primary producer, e.g. *Chaetoceros*) over the 1800 years of the ACR. Pre-ACR (18024.5 – 14619.4 cal yrs BP) the total accumulated CO_2_ of *Phaeocystis antarctica* would have been 69.19 ppm, for sea ice diatoms like *Fragilariopsis* it would have been 34.73 ppm. The modern primary producer community from the Western Antarctic Peninsula, with a dominance of *Chaetoceros* would have the best fitting accumulation of 14.13 ppm CO_2_.

These results are visualized as a time series in Figure 3B. The upper panels of Figure 3B show the NPP during the ACR and the lower panels show the NPP prior to the ACR (pre ACR). The impact of NPP on global carbon dioxide concentrations shows that a change in productivity and thus community composition from the pre ACR to the ACR is the most likely scenario. We found that the accumulated CO_2_ values assuming diatom productivity during pre-ACR and a shift to *Phaeocystis* productivity best matches the expectations from the CO_2_ curves. The figures on the left (Fig. 3B) illustrate the total amount of captured CO_2_ (ppm) subtracted from the linear model to reconstruct the actual CO_2_ trend. On the right side, the total amount of stored CO_2_ (ppm) is added to the actual CO_2_ advancement to verify whether it aligns with the linear model. A paired t-test confirmed that there are no significant differences between the captured CO_2_ by *Phaeocystis antarctica* and the actual CO_2_ progression or the linear model (p-value = 0.2219) for the ACR. However, we found significant differences of diatoms from the Ross Sea such as *Fragilariopsis* (*p-value* = 0.00005) and diatoms from the Western Antarctic Peninsula region such as *Chaetoceros* (*p-value* = 0.00001) and the actual CO_2_ curve as well as the linear model. This result underlines the higher likelihood for a dominance of *Phaeocystis antarctica* resulting in a higher drawdown of CO_2_ during the ACR compared to a diatom-dominated ecosystem.

By utilizing paleo-proxies, statistical modeling, and projection to assess the influence of net primary productions of different primary producers on global CO_2_ concentrations during the Antarctic Cold Reversal, our findings strongly indicate that *Phaeocystis antarctica*, as the dominant species, played a significant role in the heightened CO_2_ removal and potential carbon export. Our results additionally strongly suggest that no other phytoplankton species can be implicated for these observed effects. This aligns with studies of the modern Southern Ocean ecosystem indicating that carbon drawdown of *Phaeocystis antarctica* is more rapid and efficient compared to other groups of phytoplankton (9, 17).

Shifts in phytoplankton composition may also affect the global sulfur cycle, as *Phaeocystis* produces more dimethyl sulfide than diatoms (38). Therefore, we also hypothesize a positive feedback of *Phaeocystis* dominance on climate through its influence on cloud formation with higher dimethyl sulfide production (39). With a disappearance of *Phaeocystis* in the high latitude regions, dimethyl sulfide production would decrease by 7.7%, which in turn, would lead to fewer clouds and thus to a warmer climate (40). Due to climate change, a loss of *Phaeocystis* could lead to a warming of the Earth’s surface by up to 0.11°C, resulting in a reduction in annual sea-ice extent of up to 6% (40).

### Ecosystem shift at the onset of warming after the ACR

Our sedaDNA record allows unprecedented insights into the past Antarctic ecosystem covering the Antarctic Cold Reversal and the Holocene. For example, for the first time, characteristic Antarctic ice fish families (Notothenioidei) and penguin taxa were detected in a sedaDNA paleorecord. Relatively irregular signals of taxa from higher trophic levels may depend on the low biomass of consumers (approximately 30%, mainly arthropods and fishes) compared to primary producers (approximately 70%) (41). The low biomass of taxa can cause a lower number of reads and thus can have a negative impact on sedaDNA consistency (42). The ACR showed a distinct ecosystem with Channichthyidae as the dominant taxa, along with indications of the Phocidae family, while *Pleuragramma (*Nototheniidae), a key prey species (43) in the modern Ross Sea, was abundant (Fig. 2). The ecosystem during the ACR resembled the modern Ross Sea shelf region with low zooplankton and top predator abundance, suggesting a less linear food web (44) (Fig.3). With the onset of Holocene warming, Nototheniidae and the cetacean family Balaenopteridae and also Delphinidae increased in abundance, while *Pygoscelis antarcticus* (chinstrap penguin) expanded only in the Late Holocene due to the primary producer shift (45) (Fig.4). It is known that the main prey of chinstrap penguins, Antarctic krill, prefers diatoms over *Phaeocystis* (46). Additionally, the establishment of the penguin population in the Antarctic Peninsula region occurred during the Late Holocene, and was probably favored by ice-free nesting sites and (47) reduced sea-ice cover (11). The limited abundance of macro-zooplankton (Euphausiids, Iphimediidae, Lysianassidae, Temoridae, Metridinidae) and benthic taxa in sedaDNA studies could be attributed to grazing and database limitations, respectively.

## Implications and conclusion

With our study we show that sedimentary ancient DNA enables the reconstruction of past Antarctic communities of hitherto unseen completeness and taxonomic resolution. It represents the first metagenomic record of the whole ecosystem including characteristic Antarctic taxa such as *Phaeocystis antarctica*, penguins, whales, and Antarctic fishes indicating the high potential of this new proxy for future paleoecological studies (Fig. 3). Also for the first time, we were able to identify phytoplankton composition of past blooming events with the latter also being traced by the ancient bacterioplankton community record. This study shows that productivity proxies, which exclude major photoautotrophic groups like *Phaeocystis*, are only of limited use.

We identified major *Phaeocystis* blooming events and related increased carbon drawdown during a period of high seasonal sea-ice variability in the Southern Ocean during the ACR with an atmospheric CO_2_ plateau (Figs 2,3). This provides the hitherto missing piece of evidence to identify past Antarctic phytoplankton productivity as a major negative feedback in the global carbon cycle during a period of global warming. As an analogy for ongoing warming, it highlights the importance of regions with high seasonal sea-ice variability and *Phaeocystis*-dominance, such as the Ross Sea, to stabilize the atmospheric CO_2_ content. Vice versa, our results also indicate that ongoing sea-ice loss in this particular region will amplify warming because no areas farther south can take over the marine carbon pump function because of the proximity to the Antarctic continent.

Based on our reasoning, we anticipate that simulations using models parameterized/validated with diatom productivity observations likely underestimate the Antarctic carbon pump function of sea-ice impacted regions (48). However, our finding of a tremendous haptophyte-to-diatom turnover in the course of temperature increase shows how sensitive this ecosystem is to warming, potentially representing a key tipping element in the global carbon cycle. In addition, the impact of *Phaeocystis*-dominated communities on the global sulfur cycle, and thus on climate, should be given greater consideration in order to understand the effects of warming-induced changes in marine ecosystems.

As species undergo permanent adaptation it may well be possible that *Phaeocystis antarctica* decreases in abundance when adapting to a future open ocean characterized by more rare blooming conditions. And the biological function would be irreversibly lost even if physical conditions for *Phaeocystis* blooming could be reestablished.

Our study indicates the sensitivity of Antarctic ecosystems to sea-ice change (Fig. 4,5). More than that, our study suggests a complex relationship between marine fauna (e.g., penguins) and ecosystem drivers such as sea ice and temperature that points to challenges for ongoing conservation efforts.

**Figure 4:**
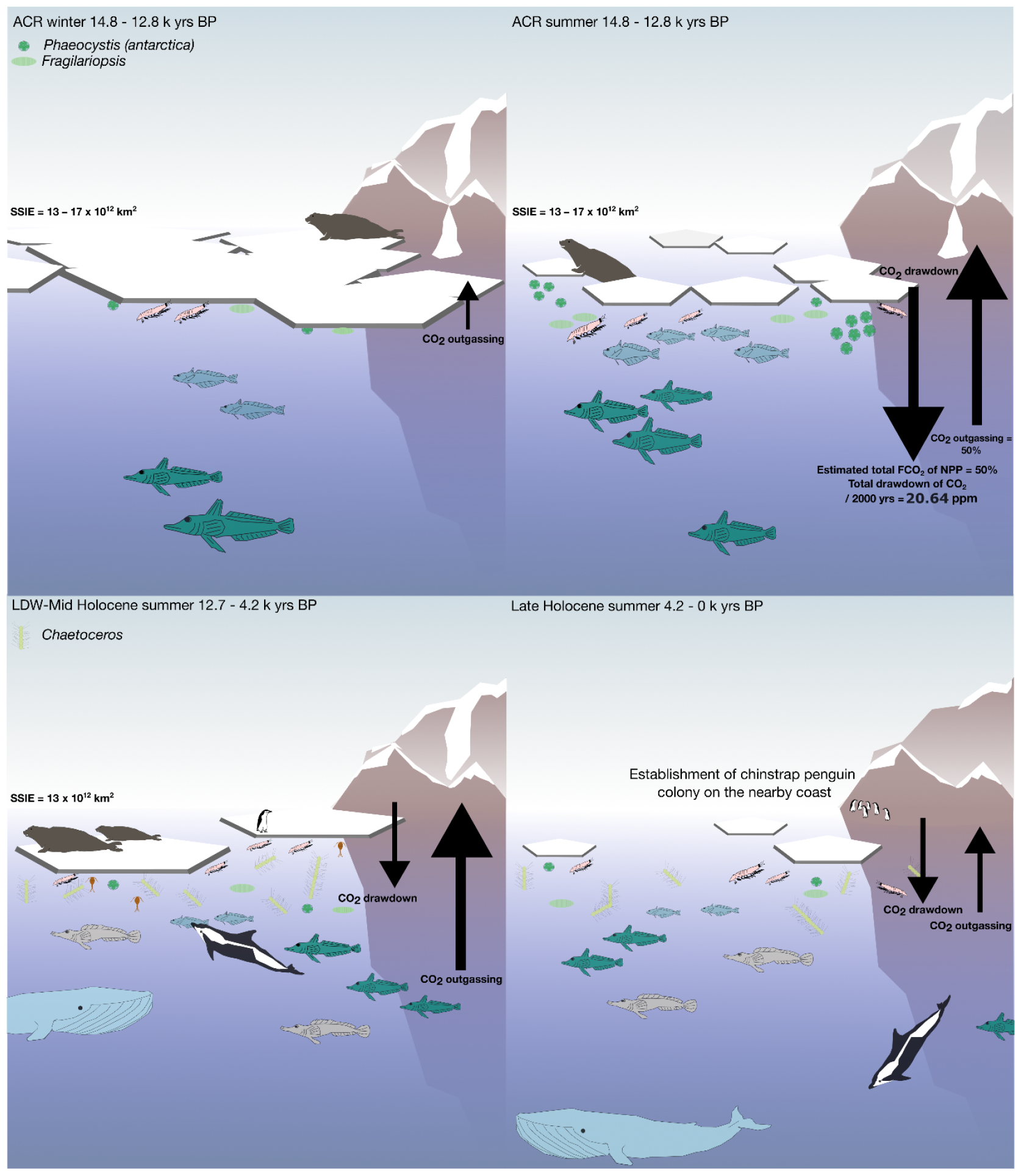
Upper panel: Seasonal ecosystem reconstruction of the Antarctic Cold Reversal inferred from sedaDNA shotgun metagenomics. The left side shows the reconstructed winter conditions in the Bransfield Strait with an indication of sea-ice coverage and glaciation of the nearby coast. The thickness of the black arrow and its direction indicate the suspected strength of outgassing of CO_2_ (8). On the right side the reconstructed ecosystem of the summer during the ACR is depicted. The black arrow, depending on its direction, indicates either the strength of outgassing or the strength of carbon drawdown. **Lower panel: Summer ecosystem reconstruction of the Late Deglacial Warming until the Middle Holocene and Middle Holocene until the Late Holocene inferred from sedaDNA shotgun metagenomics**. The left side shows the speculated ecosystem of the Bransfield Strait during the LDW, Early Holocene, and Middle Holocene. The black arrow, depending on its direction, indicates either the strength of outgassing or the strength of carbon drawdown. On the right side, the ecosystem community reconstructed for the Late Holocene is shown.

## Material and Methods

### Sediment core PS97/072-01

During the RV *Polarstern* cruise PS97 to the Drake Passage in 2016, sediment core PS97/072-1 was retrieved with a Piston corer from the east Bransfield Strait (northern Antarctic Peninsula; -62.0065, -56.0643). The core was recovered from a water depth of 1992.9 m and the total usable length was 1012 cm (51). The sediment of the core is described as a silty diatomaceous clay and the age model as well as microfossils, TOC, biogenic opal, and the biomarker data have been published by Vorrath et al. (2023) (11).

### X-ray fluorescence core scanning (XRF-CS)

Sediment core PS97/072-01 was scanned using an AVAATECH-XRF-CS. Scanning resolution was 1 cm with a detected area of 10x12 mm along the core. The elements Fe and Ba were measured with 10 kV and a 150 μA No-filter (10 s) and with 50 kV and a 1000 μA Cu-filter (20 s), respectively.

The obtained results are estimations of relative element concentrations, provided in total counts per second (cps), and are considered semi-quantitative. These estimations were derived by evaluating the intensities of the detected peak areas, as described by ref. (52). The Ba and Fe count data were normalized based on the Ba/Fe element ratio, a method established by ref. (32) (Supplementary data S3). Furthermore, the ratio was subjected to a smoothing process to enhance its consistency.

### Extraction of sedimentary ancient DNA

Halved sediment cores were subsampled under clean conditions in the climate chamber at GFZ devoid of any molecular biological work (see details in the Supplementary material). The sediment core was subsampled for 63 samples. DNA extraction of sediment samples (5–10 g wet weight) was accomplished using the *DNeasy PowerMax Soil Kit* (Qiagen, Hilden, Germany). Samples were extracted in 7 batches of nine samples and one extraction blank, which was included for generally monitoring the cleanliness of the extraction procedure.

DNA extracts were purified and concentrated with the GeneJET PCR Purification Kit (ThermoFisher Scientific, USA), DNA concentrations were measured with Qubit 4.0 fluorometer using the BR Assay Kit (Invitrogen, Carlsbad, CA, USA), and adjusted to a final concentration of 3 ng/μL.

### Shotgun metagenomics

For the metagenomic approach, 34 samples (about every ∼250 years) and 10 blanks were selected (Supplementary data 1). Associated extraction blanks were included in our six library batches, each containing 5 or 6 samples. Additionally, library blanks containing only library chemicals were added to each library batch (Supplementary data 2). Single-stranded DNA libraries were produced following the protocol of ref. (53) with modifications described in ref. (54). In short, after the library build, the libraries were quantified using qPCR to calculate the number of amplification cycles for index PCR. Sample libraries were amplified with 11–13 cycles (blank libraries with 11 cycles) using the indexing primers P5 (5’-3’: AAT GAT ACG GCG ACC ACC GAG ATC TAC ACN NNN NNN ACA CTC TTT CCC TAC ACG ACG CTC TT; IDT, Germany) and P7 (5’-3’: CAA GCA GAA GAC GGC ATA CGA GAT NNN NNN NGT GAC TGG AGT TCA GAC GTGT, IDT, Germany), the AccuPrime Pfx polymerase (Life Technologies, Germany), and 24 μL of each library. Finally, the sample libraries were pooled equimolar, while extraction blanks and library blanks were pooled with 1 μL. The library pool was purified and adjusted to the required volume (10 μL) and molarity (20nM) by using the MinElute PCR Purification Kit (Qiagen, Germany). Sequencing was conducted on a NextSeq 2000 (illumina) device using the NextSeq 2000 P3 Reagents (200 Cycles).

### Bioinformatic processing

Raw sequencing data for samples and blanks were quality checked with FastQC (version 0.11.9), and adapter trimmed and merged using Fastp (version 0.20.1). Results of the quality check of the sequencing data are given in Supplementary data S1. The taxonomic classification of passed reads was conducted with *Kraken2* (55) at a confidence threshold of 0.8 against the non-redundant nucleotide database (built with Kraken2 in April 2021).

The *Kraken2*-report files were then processed with a Python script (https://github.com/v-dinkel/downloadTaxonomicLineage) to add the full taxonomic lineage to each TaxID. Further analyses were performed in RStudio (v.1.2.5001; R v.4.0.3). Using the *dplyr* package (v.0.7.8.), the metadata, the TaxID table, and the Kraken report were merged using the *left_join* function (56).

### Ancient damage pattern analysis

The selection of key taxa for ancient damage patterns includes *Phaeocystis antarctica* (haptophyte), *Chaetoceros simplex* (diatom), *Methylophaga nitratireducenticrescens* (bacteria), and *Pseudochaenichthys georgianus* (crocodile ice fish). Prior to the analysis of damage patterns we grouped samples into five groups of adjacent samples: group 1 (0.01–7.8 k yr), group 2 (8.2–9.7 k yr), group 3 (10.5–11.4 k yr), group 4 (11.9–12.6 k yr), and group 5 (12.8–13.9 k yr). We then summed up the raw sequencing files with the *cat* command and repeated the bioinformatic processing (quality filtering, merging and classification with *Kraken2*) on the groups. The *kraken* output files were used to extract taxon-specific reads which were subsequently used for mapping reads against selected reference sequences in MapDamage (v. 2.0.8) (57) with the options rescale (to downscale quality score for mis-incorporations likely due to ancient DNA damage) and single-stranded (for a single-stranded protocol). Fragment incorporation plots and distribution of the C to T changes are presented in the Supplementary material (3. Post-mortem damage patterns).

### Statistics

The following analyses were all performed using RStudio (v.1.2.5001; R v.4.0.3). The graphics were exported from R as vector graphics (e.g., PDF) to be further enhanced with the graphics program Inkscape (Inkscape 1.1.2 (b8e25be8, 2022-02-05)). The map was created using Qgis and the Quantarctica package (v3.2) (Norwegian Polar Institute).

The data from elemental scanning were normalized using the package *compositions* (58). For the Ba/Fe ratio we used the additive logarithmic ratio transformation function (alr). To fit the Ba/Fe ratio and biomarker data to the shotgun sequencing age data points, we used the linear interpolation function (*=FORECAST*) in Excel (v. 16.74) and the interpolated data points later on for further analysis.

The EPICA Dome C (EDC) data derived from ref. (29) & ref. (50) were transformed using a translator of 650 for the secondary axis to plot both within one graphic with a 0.99 confidence level and a span of 0.1.

For the primary producer community, we only included taxa with a relative abundance greater than 1% per sample and a minimum taxonomic classification to genus level. We followed Stefan Kruse’s Github script for resampling to normalize this dataset (27) (https://github.com/StefanKruse/R_Rarefaction). For the higher trophic levels, we included only Antarctic taxa without a minimum relative abundance threshold. The relative abundance of the methylotrophic bacteria was calculated against the primary producer abundance.

The community composition over time is depicted in a stratigraphic diagram with the information of relative abundance using the R packages *tidyverse* (54), *tidypaleo* (59), and *ggplot2* (60). The arrangement of the plots was done using the *cowplot* package (61), followed by Inkscape to align them. The ecosystem drawings were done using Inkscape (Inkscape 1.1.2 (b8e25be8, 2022-02-05)).

To analyze the community and ecological turnover over time we performed a Latent Dirichlet Allocation and added a Change-point model (Supplementary Material 1). For further multivariate statistical analysis of the environmental and paleogenetic data we applied a Structural Equation Model (SEM). First we interpolated the data at consistent 300-year intervals using the rioja package *interp*.*dataset* function (62) and log transformed the phytoplankton relative abundance and temperature data. We focused on the most abundant phytoplankton taxa, including *Chaetoceros, Fragilariopsis*, and *Phaeocystis*. To uncover carbon-cycle dynamics, we used linear mixed-effects models (LMEs) with environmental factors and phytoplankton as fixed effects and time as a random effect. We fitted these LMEs with the *lme* function from the nlme package (63), examining the influence of temporal fluctuations. The model also aided the identification of alternative paths leading to a more exploratory approach due to the absence of prior hypotheses for these paths.

Subsequently, we implemented a piecewise SEM using the piecewiseSEM R package (64) to comprehensively understand the direct and indirect effects of these variables on the carbon cycle.

The projection of the impact of the net primary production on the global carbon dioxide was done using R. The cumulative atmospheric CO_2_ uptake was calculated for specific time periods, focusing on the 3 climate conditions: pre-ACR, ACR and post-ACR. This was done by grouping the data by atmospheric CO_2_ levels and calculating the cumulative uptake for each time period to identify trends of different primary producer carbon uptake in distinct environmental conditions such as the modern Ross Sea (RSE) and the modern Western Antarctic Peninsula (WAP). To create the most accurate outcome, we reduced NPP measurements in modern Southern Ocean environments to only 30% as suggested by reference Arrigo et al., 2008 due to a lower CO_2_ partial pressure in the atmosphere. Furthermore we defined the actual flux of CO_2_ (FCO_2_) to 50% which is the mean of modern FCO_2_ values for the Ross Sea environment (65). To determine the hypothetical progression of the CO_2_ concentration in the absence of a plateau, we used a linear model that accounts for both age and carbon dioxide concentration, calibrated to the increase in CO_2_ concentration during the deglacial period (18,000 - 15,000 calendar years before present), and then applied it to the ACR.

