## Supplementary Information for "*Phaeocystis* blooms caused carbon drawdown during the Antarctic Cold Reversal from sedimentary ancient DNA"

**Contents**

1. Latent Dirichlet Allocation and Change-point model
2. Information on diatom composition, concentration and biogenic opal
3. Post-mortem damage patterns

### 1. Latent Dirichlet Allocation and Change-point model

The results of a Latent Dirichlet Allocation using variational expectation-maximization (66) show two distinct communities (Supplementary Figure 2A), one with a high *Chaetoceros* proportion and one composed of *Phaeocystis*, *Fragilariopsis,* and *Micromonas* (Supplementary Figure 2B). Furthermore, results of a Bayesian change-point model, specifically developed for ecosystem data (67, 68), which calculates the time points of greatest change between communities (69) supports the ecological turnover at the end of the Antarctic Cold Reversal and the temporal trends of the main phytoplankton groups in our data (Supplementary Figure 2C).

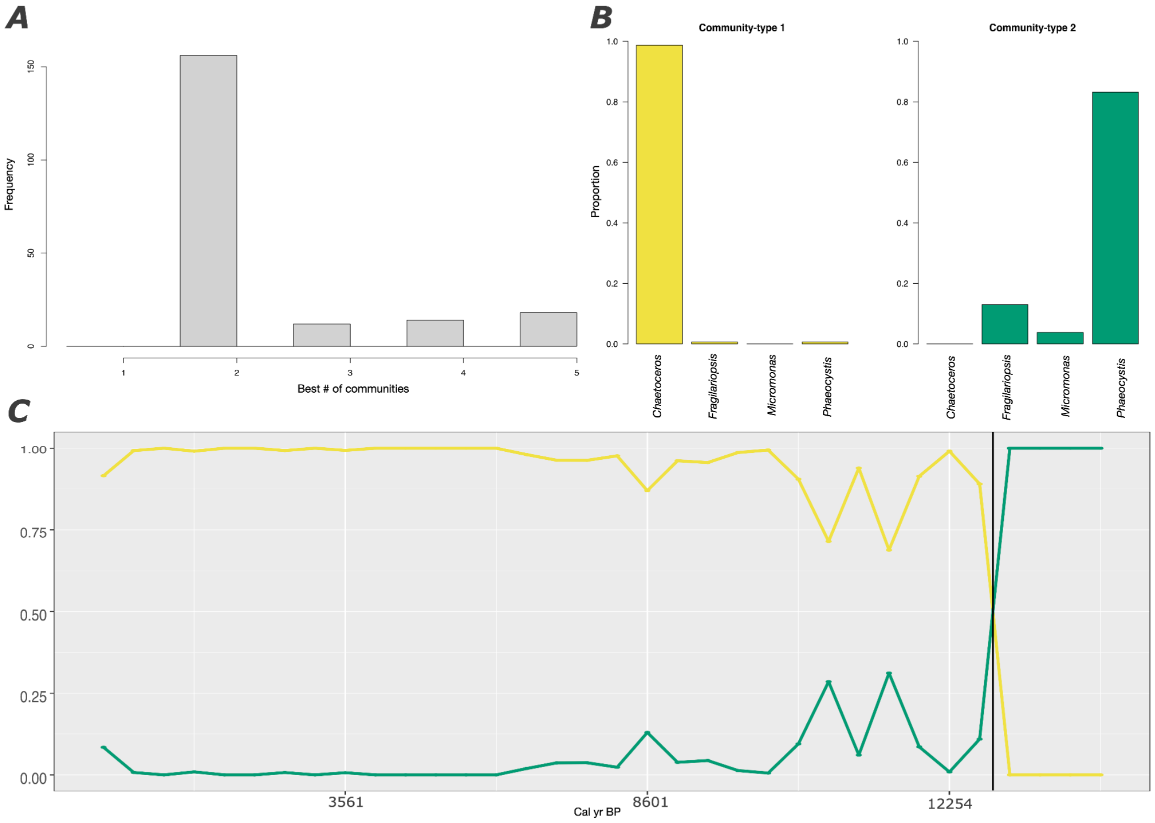


**Supplementary Figure 1: Latent Dirichlet Allocation based community analysis and change-point model analysis based on the paleogenetic community data. A** Model results of how many communities to include in the analysis. **B** Model based composition of community types 1 and 2. **C** Calculated change of the communities over time, with the black line indicating the time point of significant shift between the communities.

### 2. Information on diatom composition, concentration, and biogenic opal

The diatom composition, concentration, and biogenic opal data for the marine sediment core PS97-72/01 were previously analyzed by ref. 11 (Vorrath et al., 2023). As supplementary information, they are also included in this study. In Supplementary Figure 2, the paleogenetic data of the diatom composition are shown as well as the morphological count data of the two major diatom genera (*Chaetoceros* and *Fragilariopsis*). Biogenic opal measurements and the concentration of total diatoms are also shown.

The analysis of independent quantitative proxies reveal the same shift from low diatom abundance to high diatom abundance at the onset of the Holocene (Supplementary Figure 2). Initial biogenic opal measurements show low values during the Antarctic Cold Reversal and increase with the onset of the late deglacial warming, according to the palaeogenetic data on *Chaetoceros*, the diatom with the highest abundance after the ACR (Supplementary Figure 2). Biogenic opal may be affected by radiolarians living in symbiosis with *Phaeocystis*

(70), but there are only very few in the paleogenetic record. The Pearson correlations, calculated using the *Corit* R package (https://github.com/EarthSystemDiagnostics/corit), of both datasets show the same results with a significant positive correlation (Supplementary Table 1). Accordingly, there was not a high export of diatoms during the ACR compared to the Holocene. In addition, total diatom concentration (valve/g) is low during the ACR, increasing only between 12,800 and 11,000 cal yr BP (Supplementary Figure 2). Comparison of the qualitative community composition inferred from morphological count data and paleogenetic data reveals significant positive correlation between CRS$^{Morphology}$ and *Chaetoceros*$^{Paleogenetic}$ as well as *Fragilariopsis*$^{Morphology}$ and *Fragilariopsis*$^{Paleogenetic}$ (Supplementary Table 1).

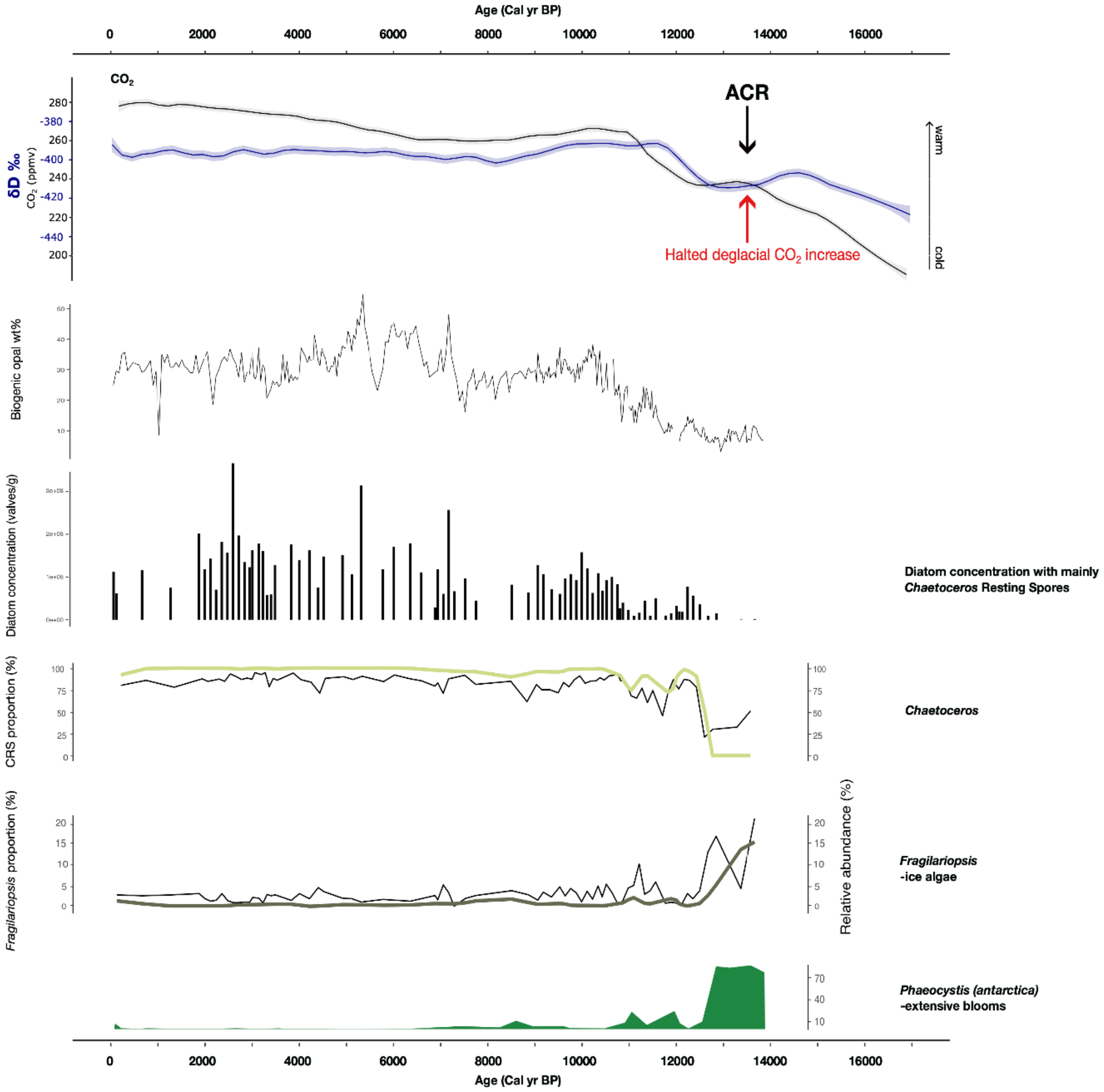


**Supplementary Figure 2: Biogenic opal (wt%) (11) and diatom concentration with CRS (Chaetoceros Resting Spores) (valves/g) (11) over time (cal kyr BP). Comparison of morphological count data and paleogenetic data of *Chaetoceros* and *Fragilariopsis*.** For comparison the $CO_2$ concentration (48) and δD ‰ (29) of EPICA Dome C (top) as well as the relative abundance of *Phaeocystis* (bottom) are shown.

**Supplementary Table 1: Pearson correlation of paleogenetic diatom data (relative abundance %), morphological composition (proportion % of Chaetoceros Resting Spores and *Fragilariopsis* valves), biogenic opal (wt%), and total valve concentration (valves/g).**

| | CRS$^{Morphology}$ (proportion %) | *Fragilariopsis*$^{Morphology}$ *(*proportion *%)* | Biogenic opal wt% | Diatom Concentration (valves/g) |
|---|---|---|---|---|
| *Chaetoceros*$^{Paleogenetic}$ (rel. abund. %) | $R$ = 0.89 p = 0.005 | - | $R$ = 0.67 p = 0.005 | $R$ = 0.54 p = 0.015 |
| *Fragilariopsis*$^{Paleogenetic}$ (rel. abund. %) | - | $R$ = 0.89 p = 0.005 | - | - |

### 3. Post-mortem damage patterns

The selection of key taxa for post-mortem damage patterns includes the haptophyte *Phaeocystis antarctica* (used reference genome for alignment: complete plastid genome JN117275.2), the diatom *Chaetoceros simplex* (used reference genome for alignment: complete plastid genome NC_025310.1), the bacteria *Methylophaga nitratireducenticrescens* (used reference genome for alignment: RefSeq assembly containing one complete genome GCF_000260985.4), and the crocodile icefish *Pseudochaenichthys georgianus* (used reference genome for alignment: RefSeq assembly containing single chromosomes GCF_902827115.2). Prior to the analysis of damage patterns we grouped samples into five groups of adjacent samples belonging to similar geo-climatic periods: **group 1** Early to Middle Holocene (0.01–7.8 k yr, 17 samples), **group 2** Late Holocene (8.2–9.7 k yr, 5 samples), **group 3** Transition phase between Holocene and Pleistocene (10.5–11.4 k yr, 4 samples), **group 4** Late Deglacial Warming (11.9–12.6 k yr, 4 samples), and **group 5** Antarctic Cold Reversal (12.8–13.9 k yr, 4 samples). The grouping of samples was done to increase the read numbers for each taxon, which improves the damage pattern analysis. Due to the absence of *Chaetoceros simplex* in age group 5, no post-mortem damage plot was generated. The post-mortem signatures are identified by an increased frequency of C to T substitutions at the ends of the DNA strands, which is identified by the comparison of reads against the relevant reference genome. The frequency of C to T changes at the end (last 25 base pairs) of the paired-end reads, the four frequency panels, showing the frequency of the four bases outside and in the read (the open gray box corresponds to the read), the posterior prediction plots, and the read length distribution plots are presented in **Figures S3-S21**. We identify higher variation in the four frequency panels for the crocodile icefish *Pseudochaenichthys georgianus*, compared to the other taxa, as a result of the multiple mappings against the single chromosomes provided in the reference RefSeq assembly used for the mapdamage analysis.

For **Figures S3-S21** panel **A** shows the four frequency graphs, displaying the frequency of the four bases outside and in the read (the open gray box corresponds to the read); frequency of C to T changes at a position's specific substitution from the 5' (left) and the 3' end (right) of the paired-end reads. Color codes are red: C to T substitutions, blue: G to A substitutions, gray: all other substitutions, orange: soft-clipped bases, purple: insertions relative to the reference. Panel **B**: shows posterior prediction plots giving the empirical misincorporation frequency and the posterior predictive intervals from the fitted model. Panel **C** shows the distribution of read length of merged reads, which are mapped to the relevant reference genomes.

**Supplementary Figure 3-21**

**A**

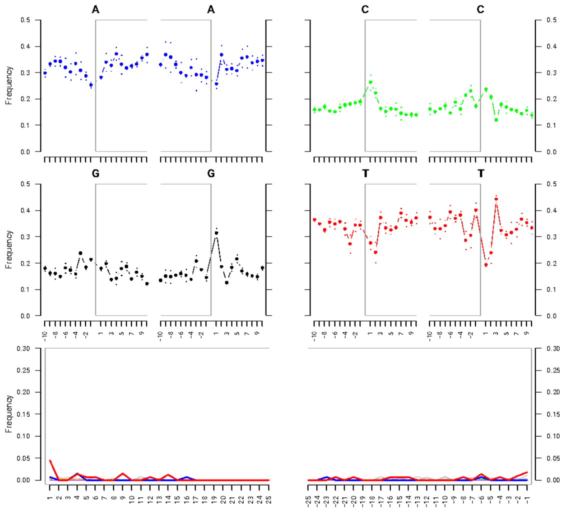


**B**

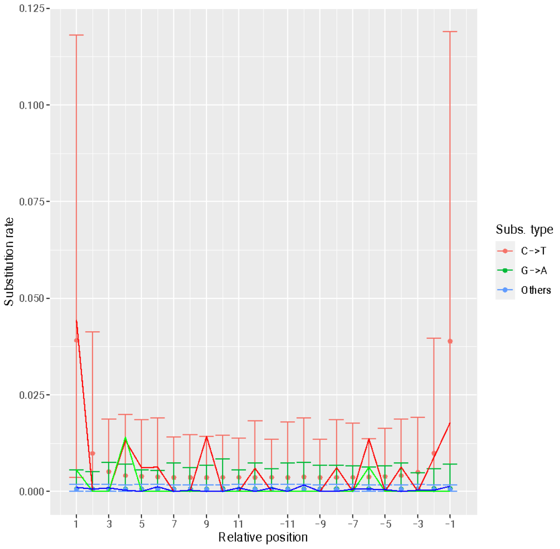


**C**

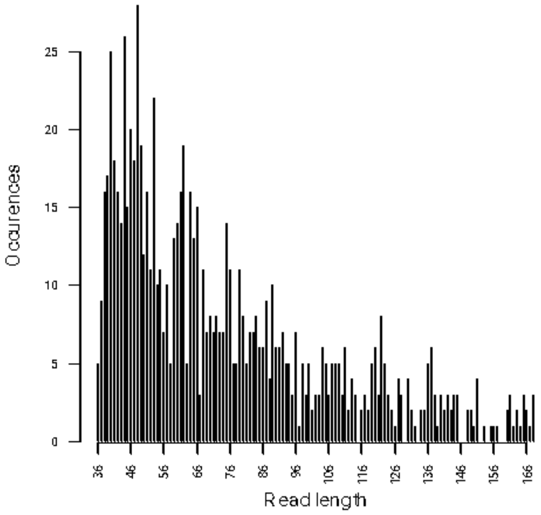


**Supplementary Figure 3:** Post-mortem patterns for *Phaeocystis antarctica* in group 1 (96–7783 cal yr BP; number of total reads extracted: 1,072, whereof 981 (91.5%) are mapped to the reference).

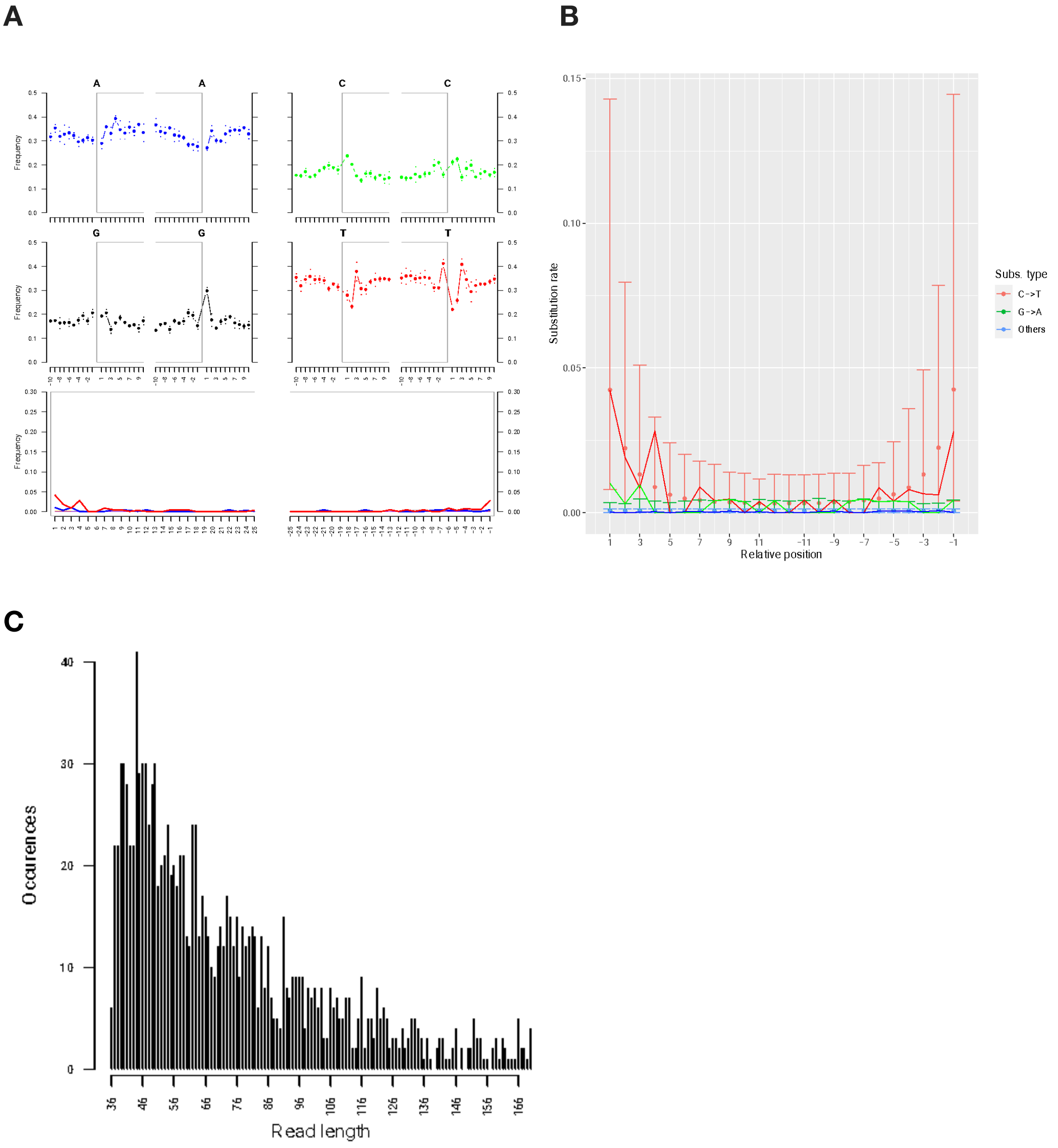


**Supplementary Figure 4:** Post-mortem patterns for *Phaeocystis antarctica* in group 2 (8250–9739 cal yr BP; number of total reads extracted: 1,358 whereof 1,277 (94%) are mapped to the reference).

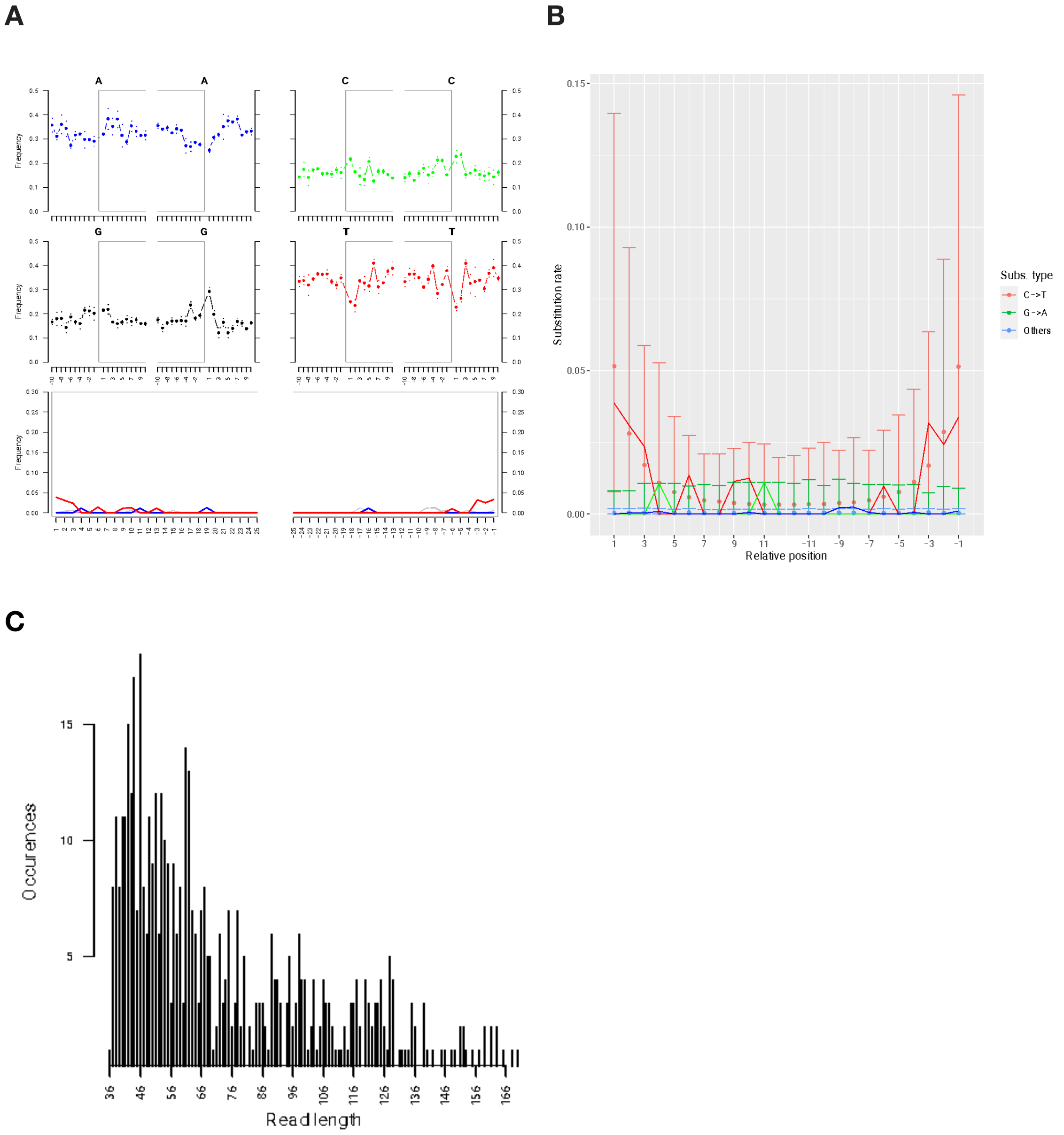


**Supplementary Figure 5:** Post-mortem patterns for *Phaeocystis antarctica* in group 3 (10469–11374 cal yr BP; number of total reads extracted: 519 whereof 505 (97.3%) are mapped to the reference).

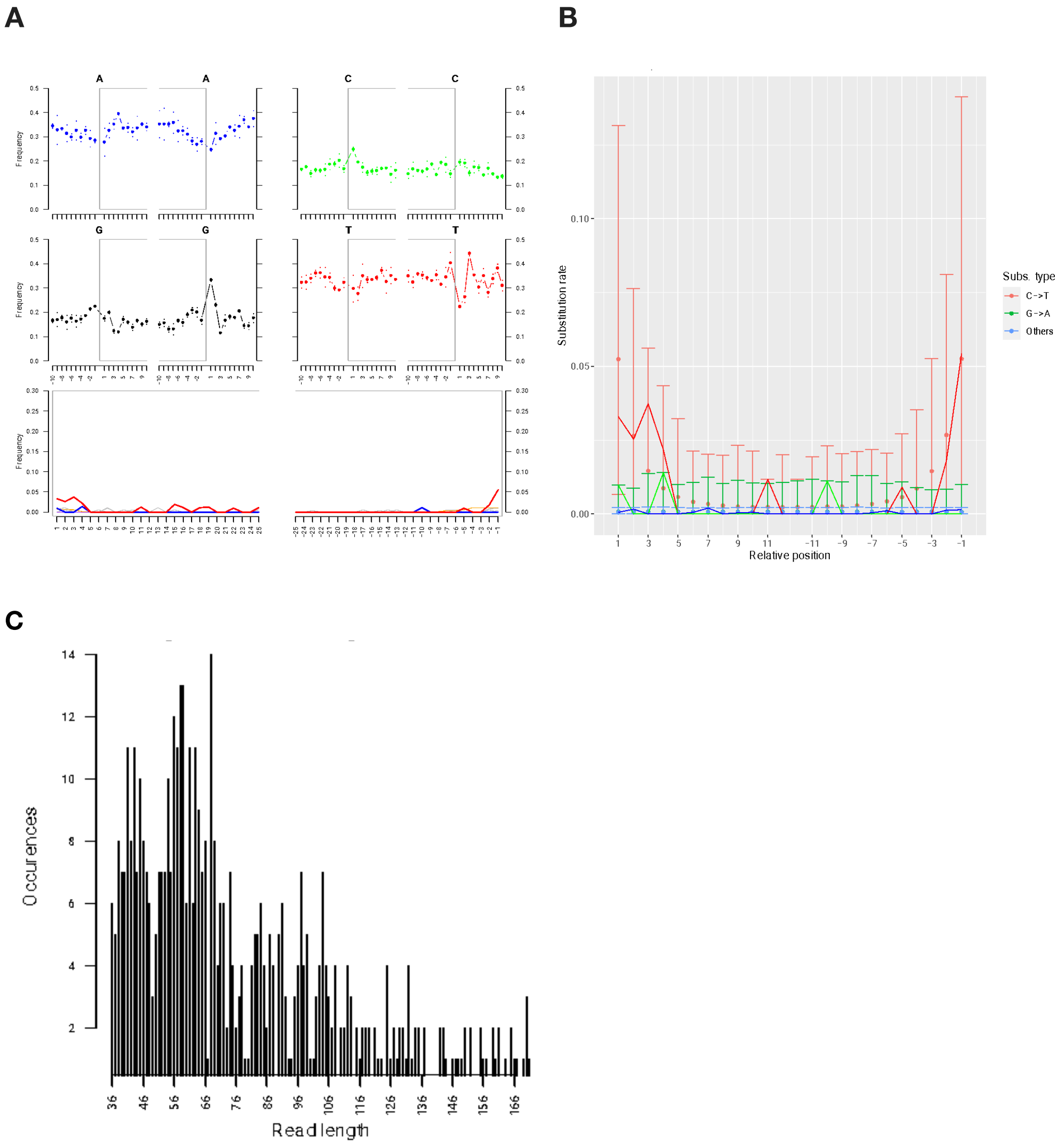


**Supplementary Figure 6:** Post-mortem patterns for *Phaeocystis antarctica* in group 4 (11960–12547 cal yr BP; number of total reads extracted: 533 whereof 500 (93.8%) are mapped to the reference).

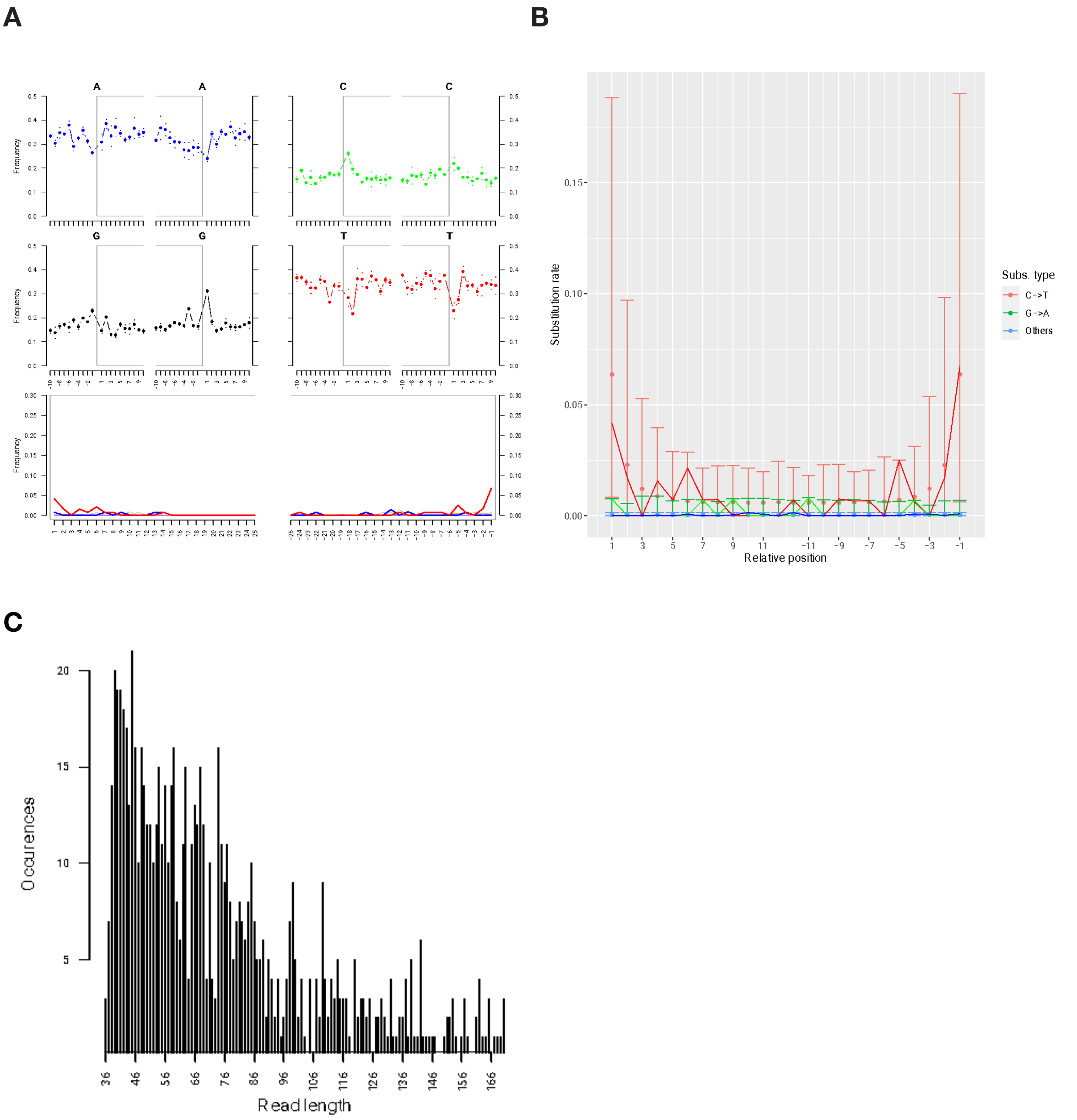


**Supplementary Figure 7:** Post-mortem patterns for *Phaeocystis antarctica* in group 5 (12840–13867 cal yr BP; number of total reads extracted: 829 whereof 781 (94.2%) are mapped to the reference).

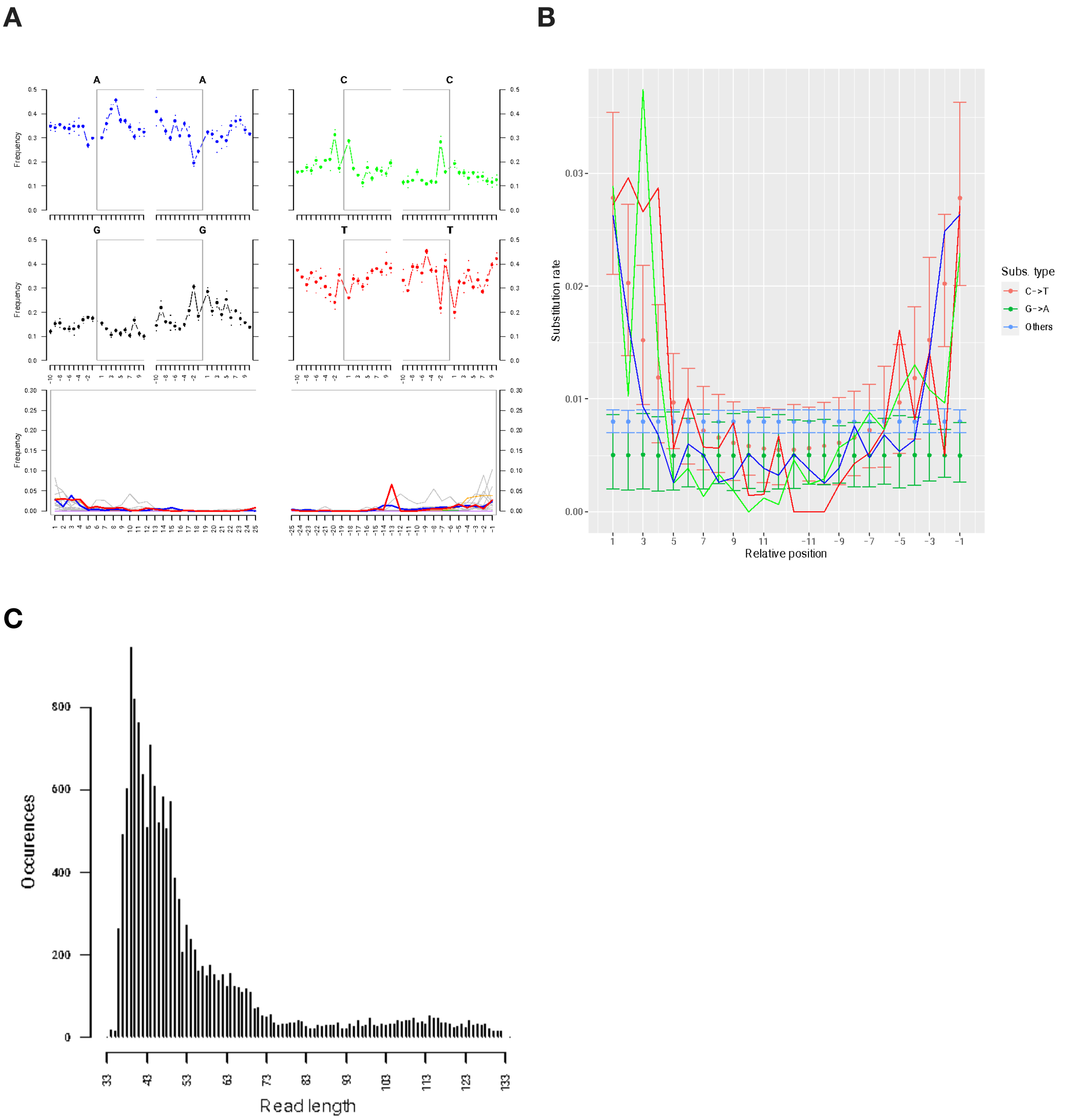


**Supplementary Figure 8:** Post-mortem patterns for *Chaetoceros simplex* in group 1 (96–7783 cal yrs BP; number of total reads extracted: 14,339 whereof 14,258 (99.4%) are mapped to the reference).

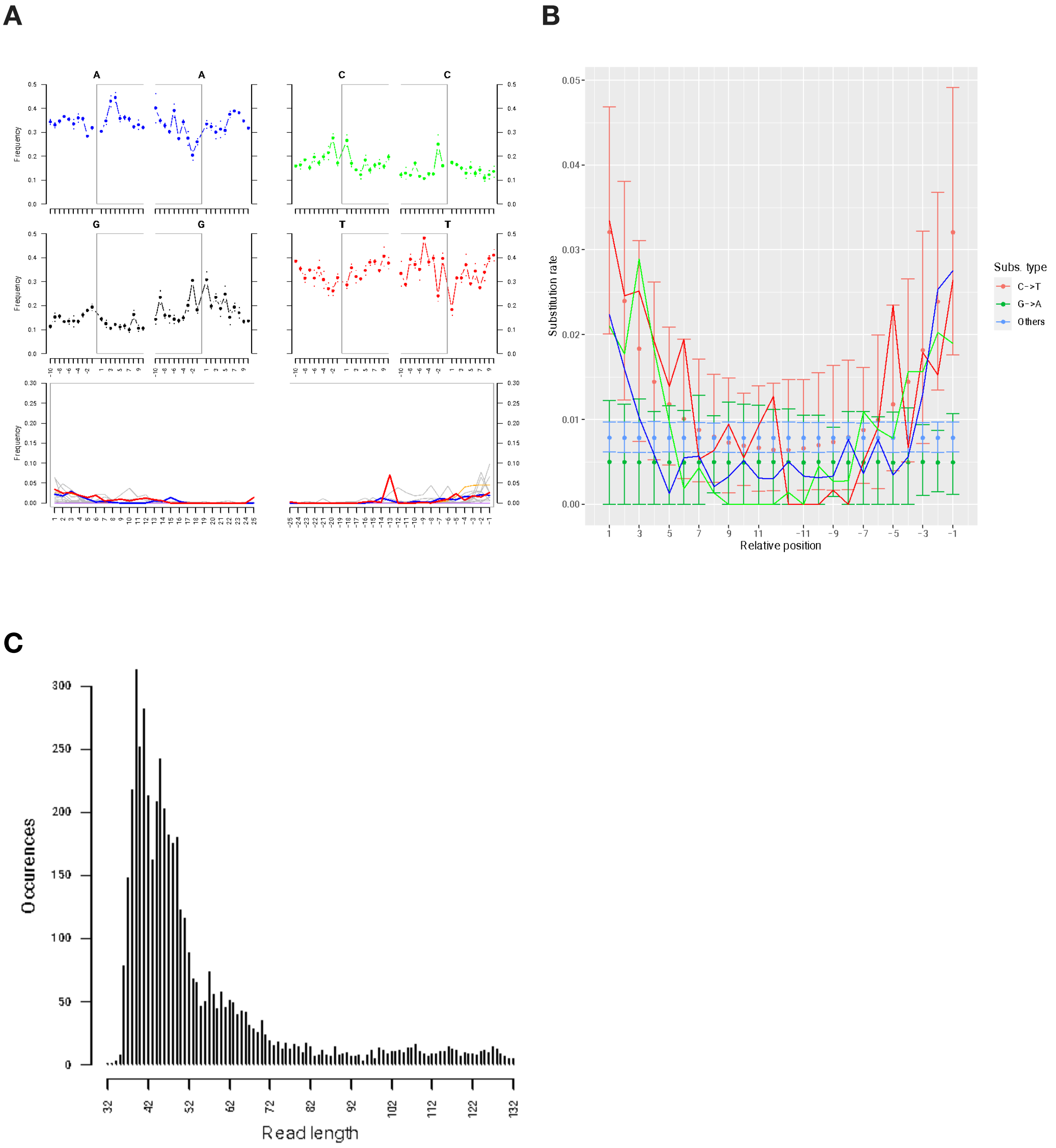


**Supplementary Figure 9:** Post-mortem patterns for *Chaetoceros simplex* in group 2 (8250–9739 cal yr BP; number of total reads extracted: 4,694 whereof 4,672 (99.5%) are mapped to the reference).

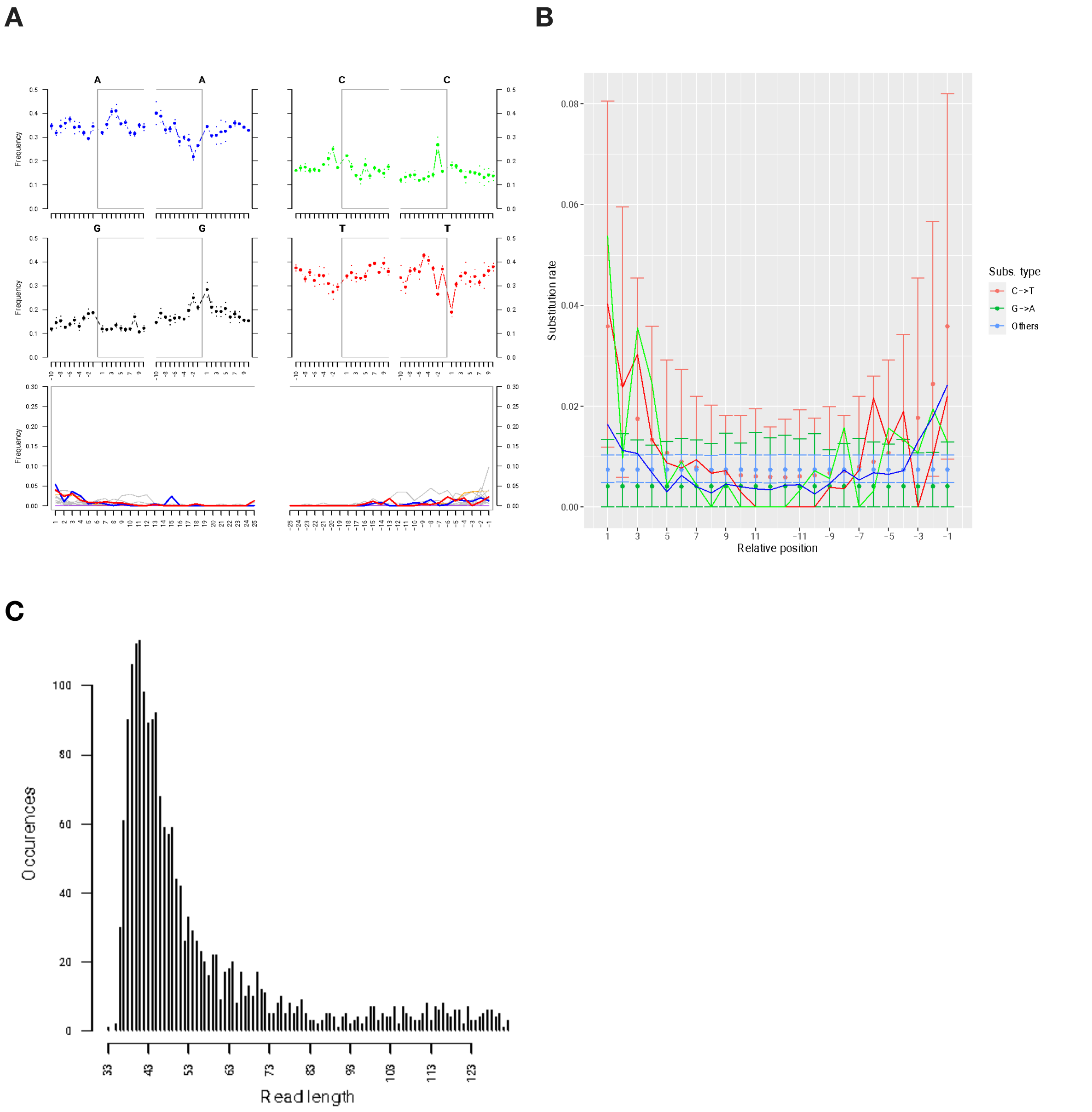


**Supplementary Figure 10:** Post-mortem patterns for *Chaetoceros simplex* in group 3 (10469–11374 cal yr BP; number of total reads extracted: 1891 whereof 1877 (99.3%) are mapped to the reference).

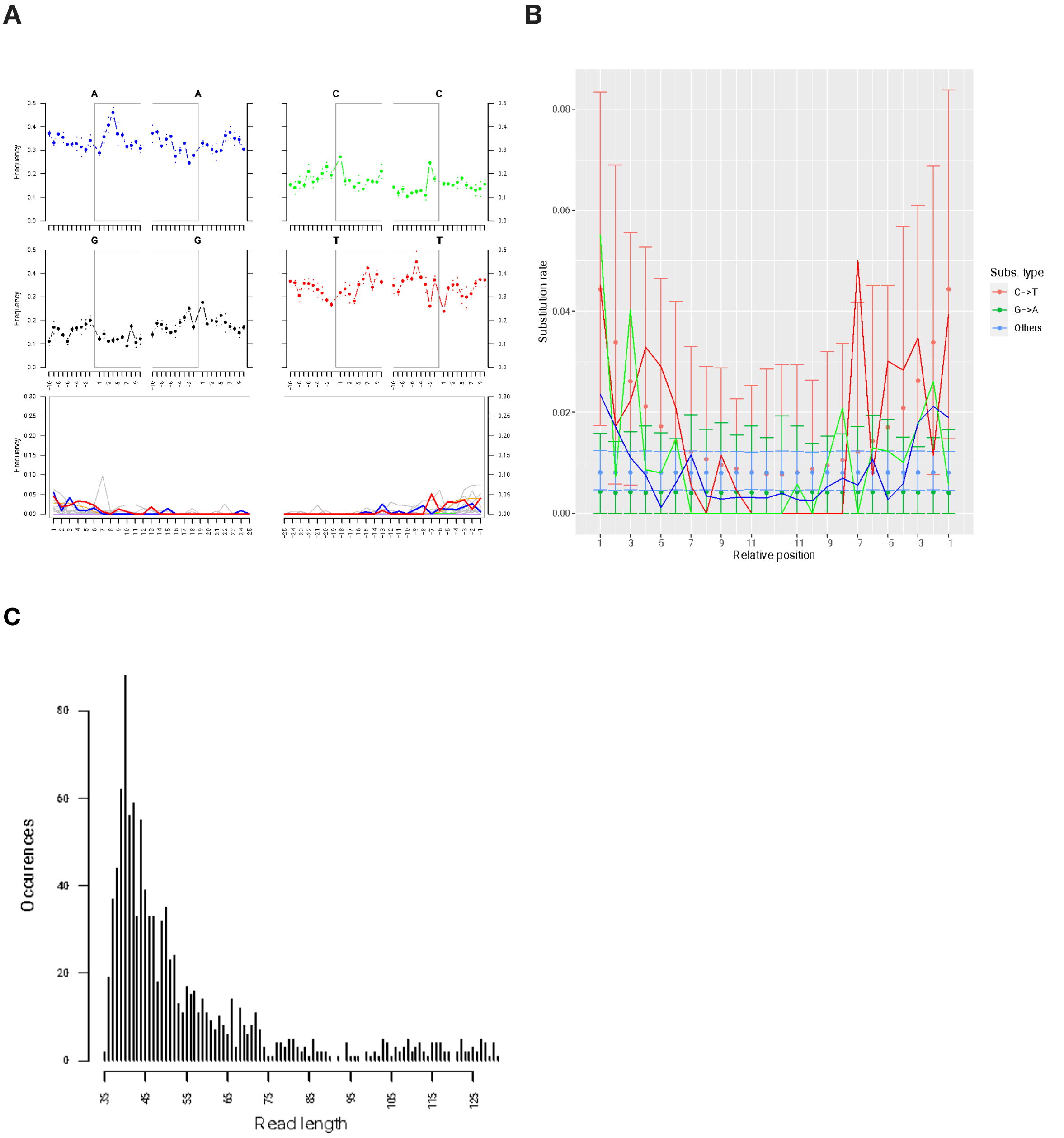


**Supplementary Figure 11:** Post-mortem patterns for *Chaetoceros simplex* in group 4 (11960–12547 cal yr BP; number of total reads extracted: 1,059 whereof 1,052 (99.3%) are mapped to the reference).

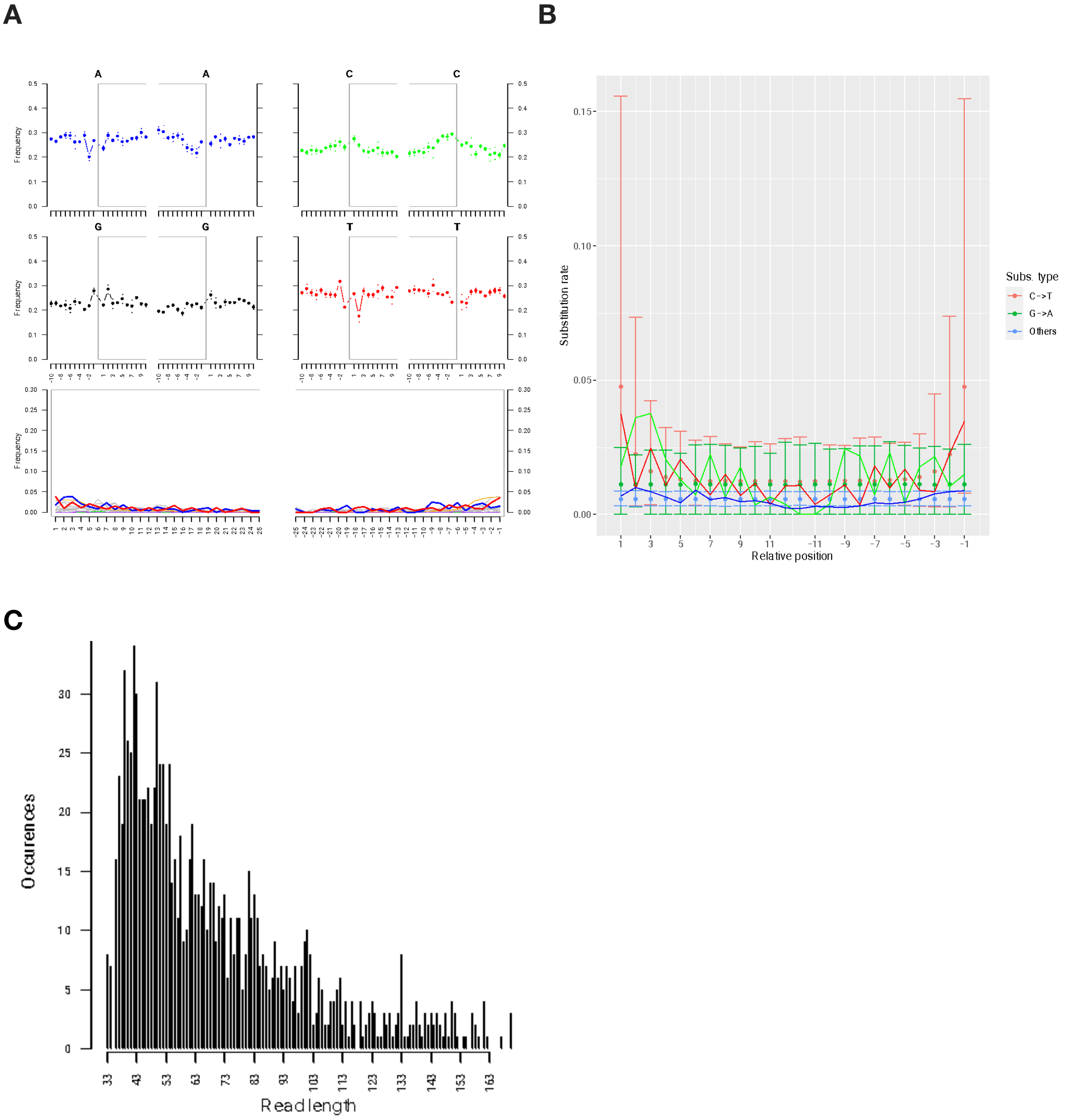


**Supplementary Figure 12:** Post-mortem patterns for *Methylophaga nitratireducenticrescens* in group 1 (96–7783 cal yr BP; number of total reads extracted: 1,311 whereof 1,253 (95.6%) are mapped to the reference).

**A**

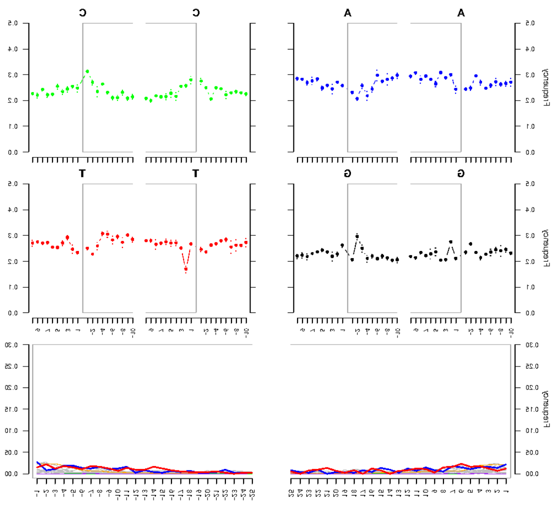

**B**

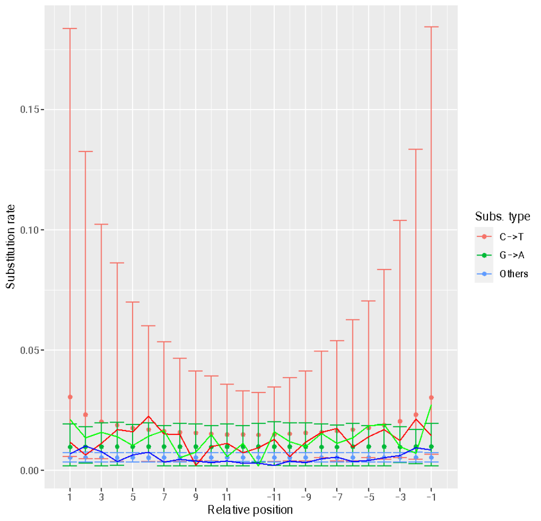


**C**

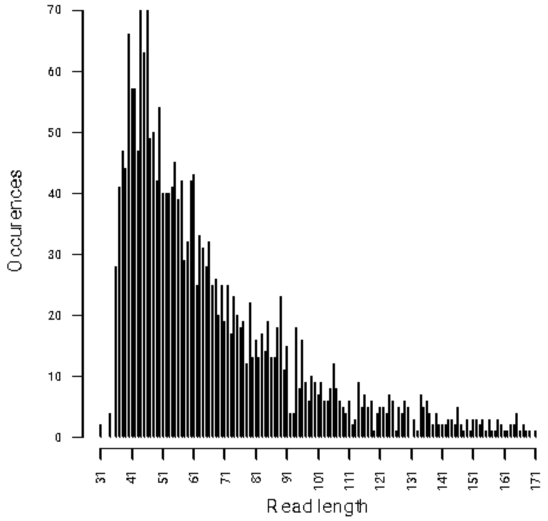


**Supplementary Figure 13:** Post-mortem patterns for *Methylophaga nitratireducenticrescens* in group 2 (8250–9739 cal yr BP; number of total reads extracted: 2,261 whereof 2,177 (95.6%) are mapped to the reference).

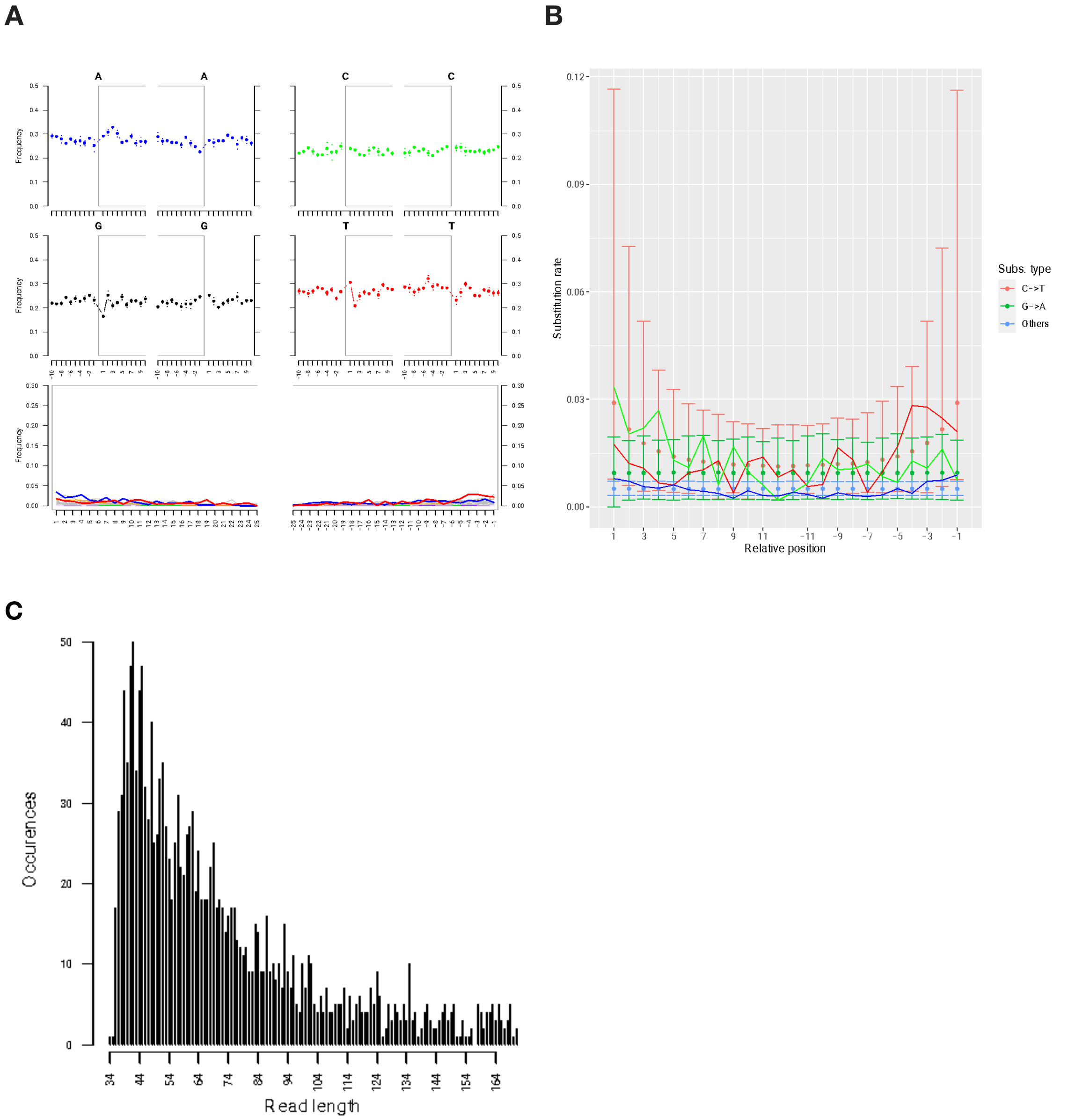


**Supplementary Figure 14:** Post-mortem patterns for *Methylophaga nitratireducenticrescens* in group 3 (10469–11374 cal yr BP; number of total reads extracted: 1,671 whereof 1,637 (98%) are mapped to the reference).

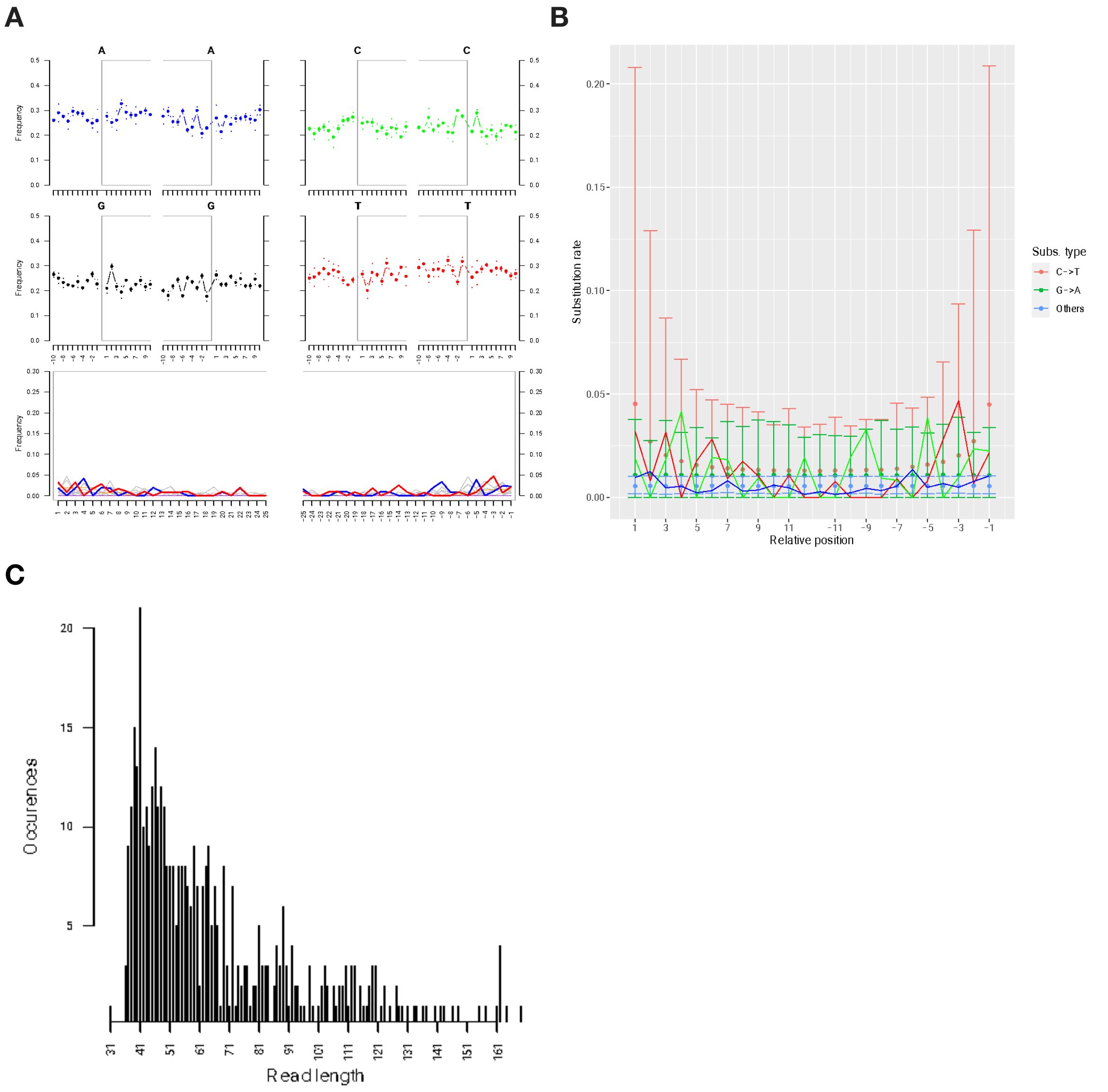


**Supplementary Figure 15:** Post-mortem patterns for *Methylophaga nitratireducenticrescens* in group 4 (11960–12547 cal yr BP; number of total reads extracted: 509 whereof 463 (85.7%) are mapped to the reference).

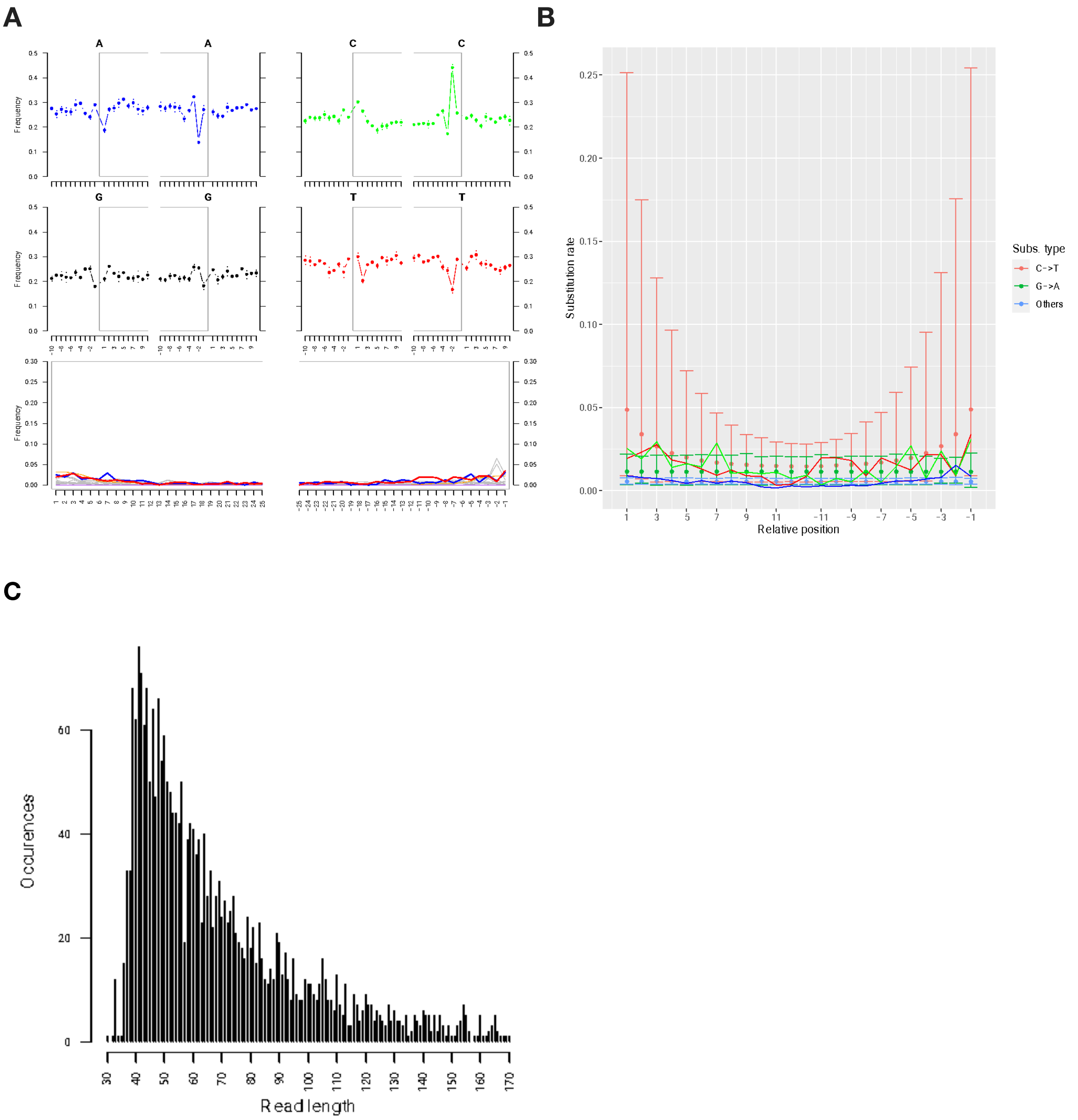


**Supplementary Figure 16:** Post-mortem patterns for *Methylophaga nitratireducenticrescens* in group 5 (12840–13867 cal yr BP; number of total reads extracted: 2,575 whereof 2,401 (93.2%) are mapped to the reference).

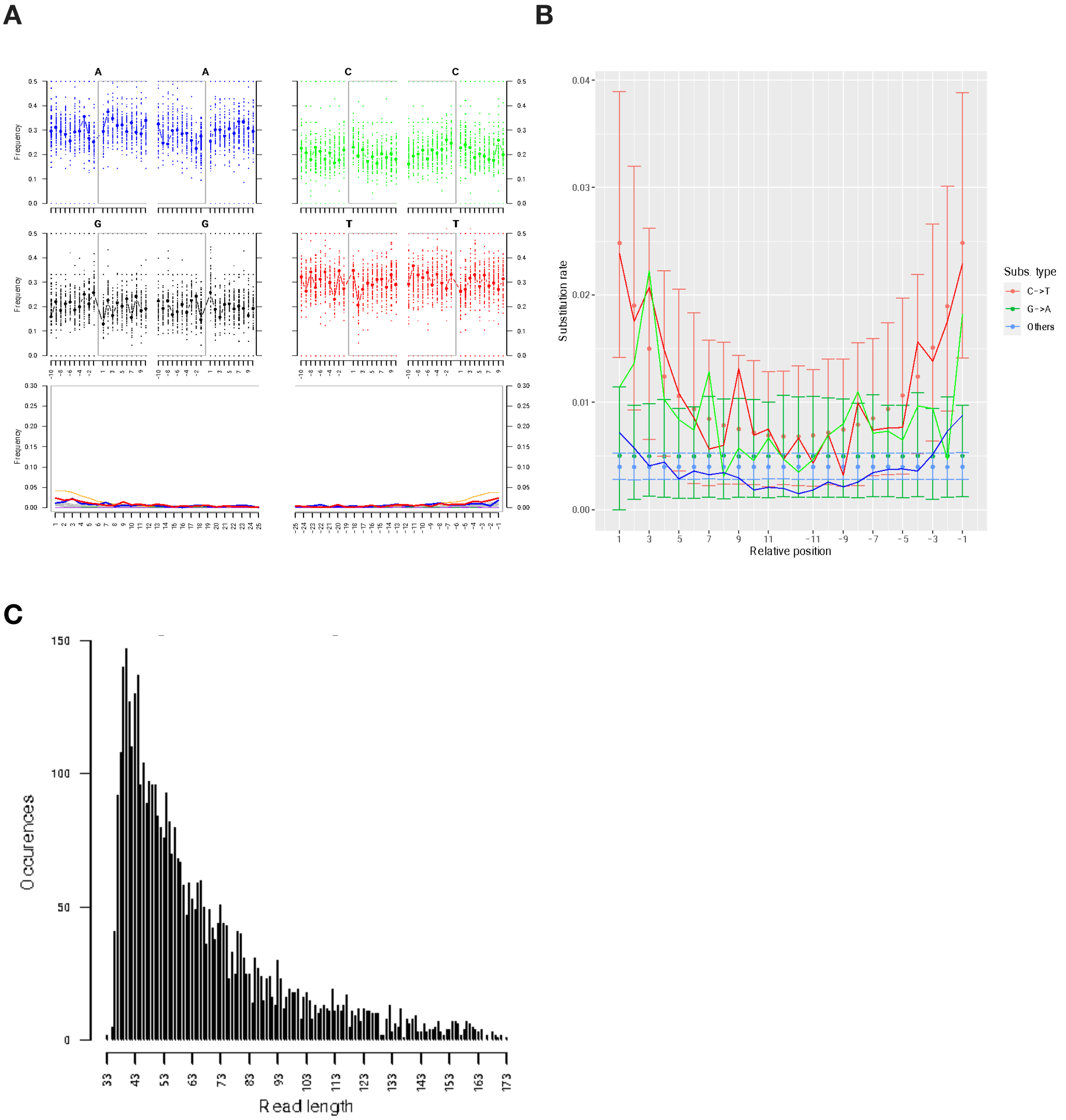


**Supplementary Figure 17:** Post-mortem patterns for *Pseudochaenichthys georgianus* in group 1 (96–7783 cal yr BP; number of total reads extracted: 4,513 whereof 4,509 (99.9%) are mapped to the reference).

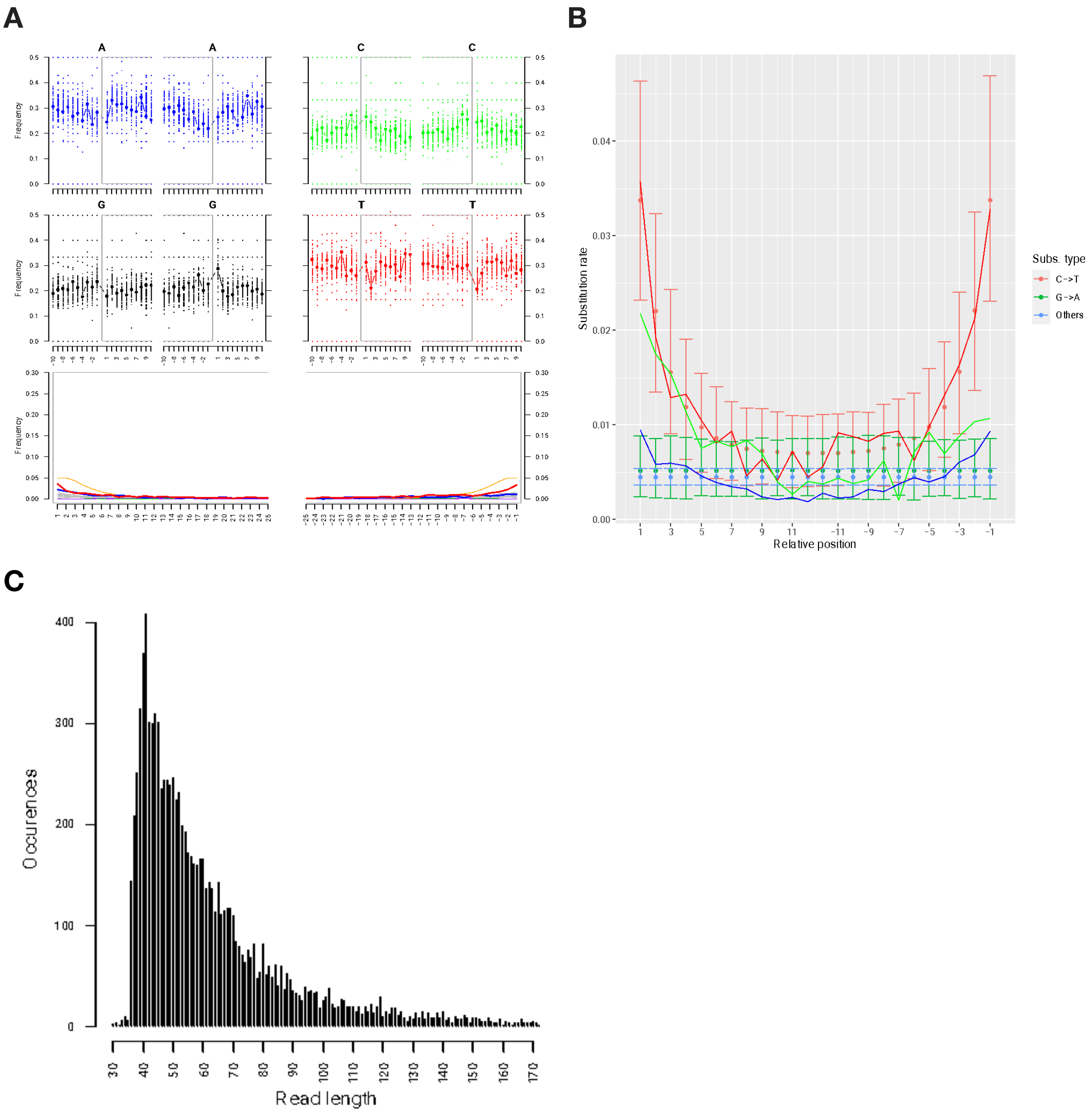


**Supplementary Figure 18:** Post-mortem patterns for *Pseudochaenichthys georgianus* in group 2 (8250–9739 cal yr BP; number of total reads extracted: 9,538 whereof 9,529 (99.9%) are mapped to the reference).

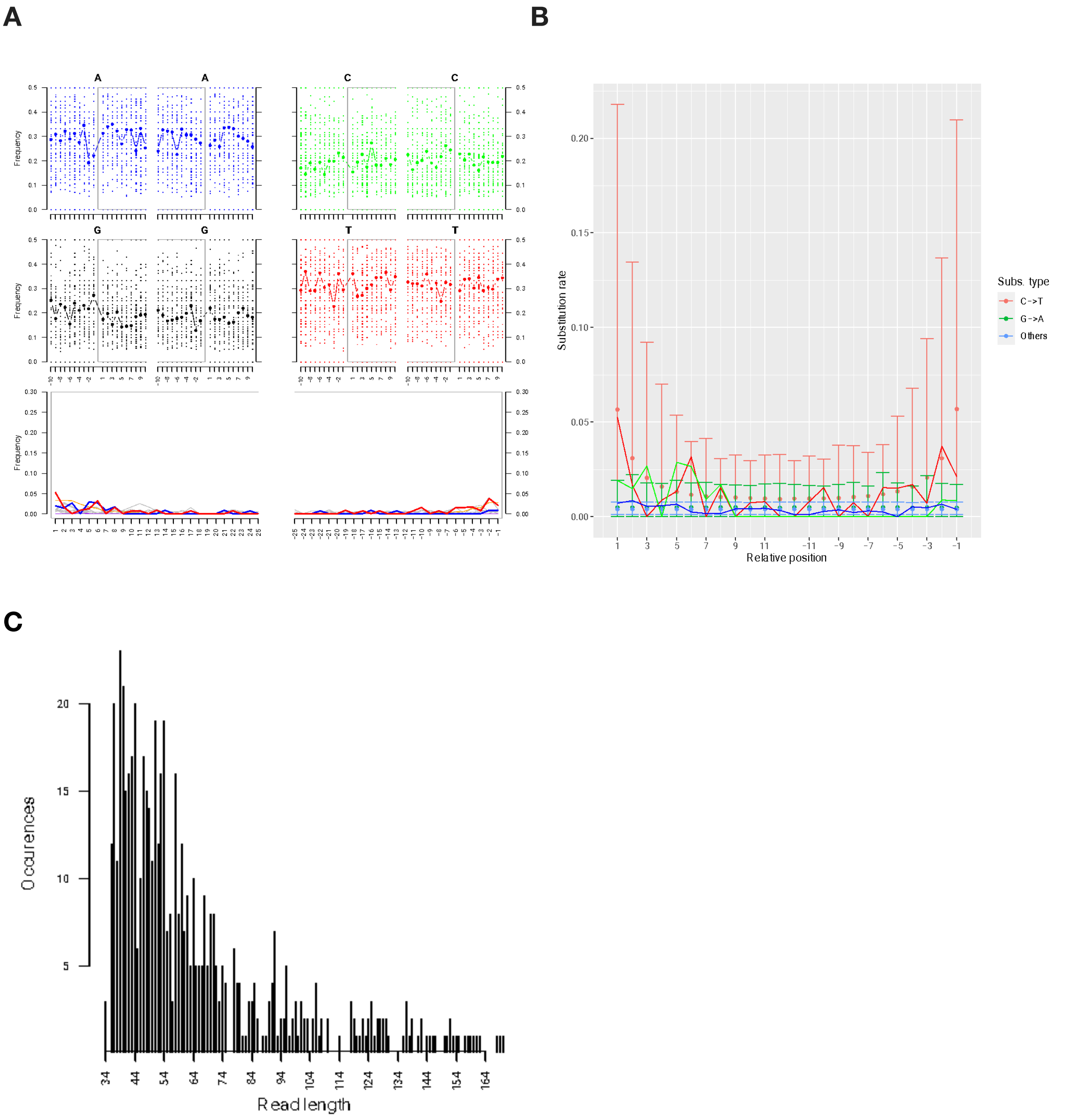


**Supplementary Figure 19:** Post-mortem patterns for *Pseudochaenichthys georgianus* in group 3 (10469–11374 cal yr BP; number of total reads extracted: 579 whereof 579 (100%) are mapped to the reference).

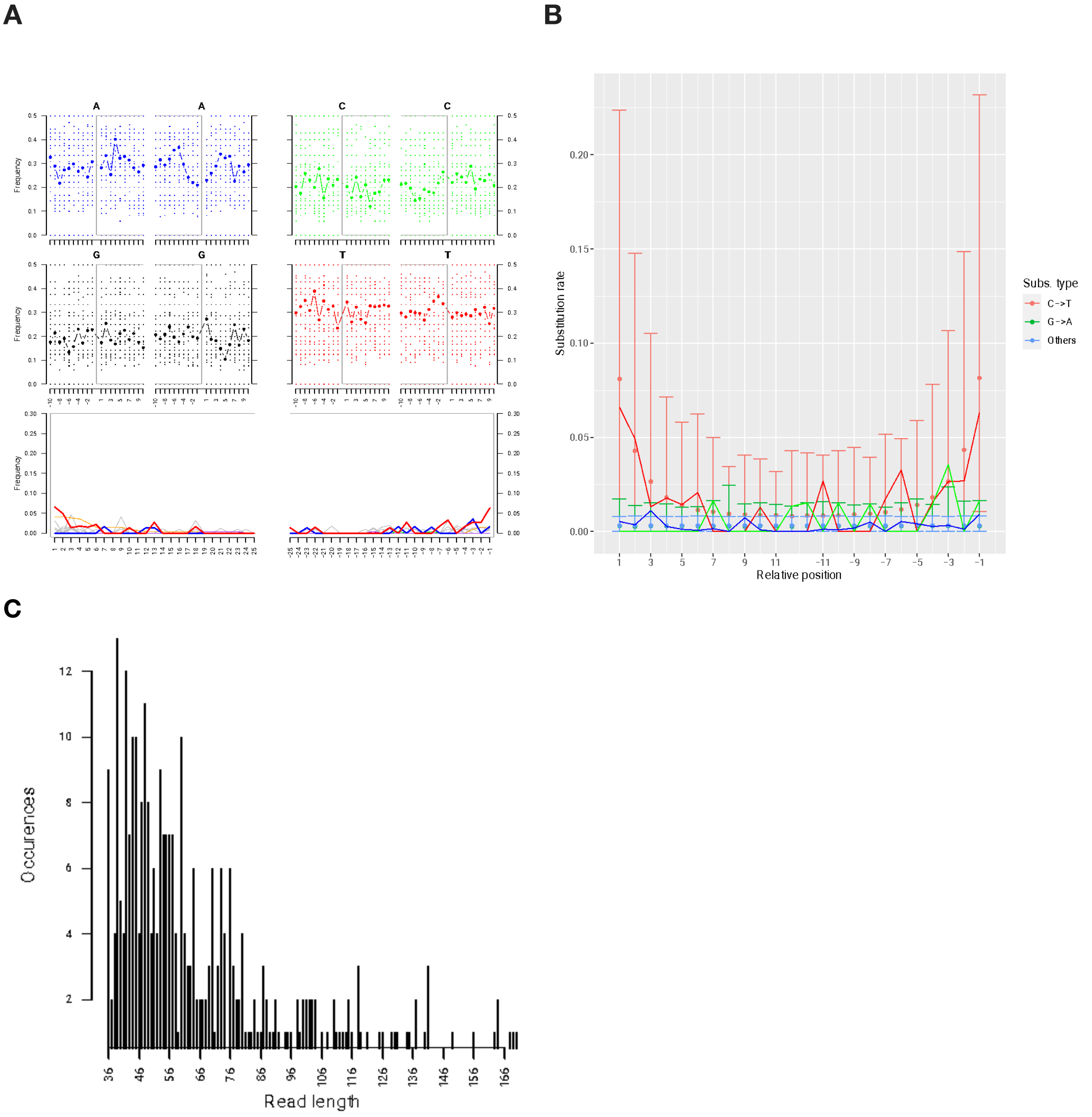


**Supplementary Figure 20:** Post-mortem patterns for *Pseudochaenichthys georgianus* in group 4 (11960–12547 cal yr BP; number of total reads extracted: 312 whereof 307 (98.4%) are mapped to the reference).

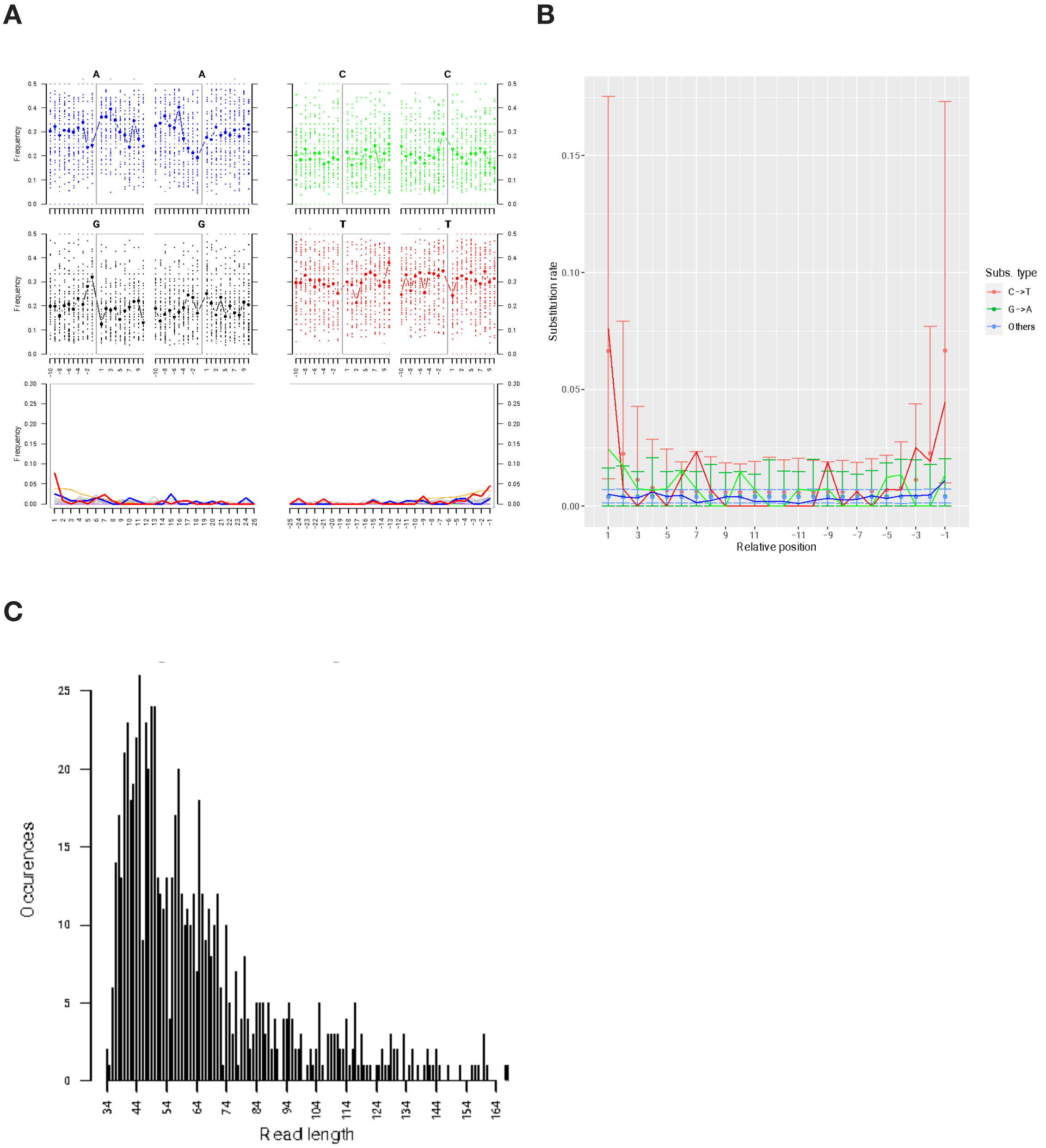


**Supplementary Figure 21:** Post-mortem patterns for *Pseudochaenichthys georgianus* in group 5 (12840–13867 cal yr BP; number of total reads extracted: 722 whereof 722 (100%) are mapped to the reference).
